# Evaluating computational efforts and physiological resolution of mathematical models of cardiac tissue

**DOI:** 10.1101/2024.02.27.582288

**Authors:** Karoline Horgmo Jæger, James D. Trotter, Xing Cai, Hermenegild Arevalo, Aslak Tveito

## Abstract

Computational techniques have significantly advanced our understanding of cardiac electrophysiology, yet they have predominantly concentrated on averaged models that do not represent the intricate dynamics near individual cardiomyocytes. Recently, accurate models representing individual cells have gained popularity, enabling analysis of the electrophysiology at the micrometer level. Here, we evaluate five mathematical models to determine their computational efficiency and physiological fidelity. Our findings reveal that cell-based models introduced in recent literature offer both efficiency and precision for simulating small tissue samples (comprising thousands of cardiomyocytes). Conversely, the traditional bidomain model and its simplified counterpart, the monodomain model, are more appropriate for larger tissue masses (encompassing millions to billions of cardiomyocytes). For simulations requiring detailed parameter variations along individual cell membranes, the EMI model emerges as the only viable choice. This model distinctively accounts for the extracellular (E), membrane (M), and intracellular (I) spaces, providing a comprehensive framework for detailed studies. Nonetheless, the EMI model’s applicability to large-scale tissues is limited by its substantial computational demands for subcellular resolution.

## 1 Introduction

Numerical simulations of cardiac cells have traditionally followed two primary approaches: membrane models, which are expressed as systems of ordinary differential equations, see, e.g., [1, 2], or spatial models, such as the bidomain model (BD) and monodomain model (MD), see, e.g., [3, 4]. Membrane models, due to their lack of spatial variation, are unable to explain important spatial phenomena like reentry, arrhythmias, or fibrillation. These spatial aspects have been extensively investigated using the BD and MD models. However, these models are based on homogenization or averaging approaches (see [3, 5, 6, 7]), leading to the removal of the cardiomyocyte from the simulations. While this simplifies the models, it also significantly restricts their resolution. As a result, BD and MD are not well-suited for modeling electrophysiological processes at the level of individual cells, which is a critical limitation in understanding cardiac dynamics and the origin of arrhythmias. With the continuous increase in computing power and advancements in numerical solution techniques for partial differential equations, the potential for enhancing the resolution of cardiac tissue models has become evident. This advancement has been a key focus of the MICROCARD project [8], which aims to provide simulation software capable of sub-cellular resolution. Such software is pivotal for analyzing the progression from cellular and sub-cellular level perturbations to the initiation of cardiac arrhythmias.

In the traditional BD and MD models, the extracellular space (E), the cell membrane (M), and the intracellular space (I) are uniformly distributed throughout the computational domain. However, more accurate cell-based models distinctly separate these domains. Consequently, these models are now often referred to as EMI models, but they were earlier referred to as microscopic models, see, e.g., [9]. The development of EMI models is presented in [10, 11, 12, 13, 14, 15, 16], and their applications are explored in [17, 18, 19, 20, 21, 22]. Numerical methods for solving the EMI equations have been analyzed in [23, 24, 25, 26, 27]. Additionally, the properties and efficiencies of these solvers have been evaluated in [28, 29, 30, 31, 7, 23, 32], and special methods for mesh generation for EMI models have been addressed in [33].

The use of EMI models is associated with a considerable increase in computational demands. The extent of the increase depends on the actual application under consideration, but from the definition of the models it is quite clear that whereas the BD and MD models provide average solutions over many cardiomyocytes, the EMI models resolve every cell into a large number of mesh blocks, see, e.g., [13]. For specific applications, it may very well occur that BD and MD are too coarse but the EMI models are more accurate than what is actually required. This has motivated the development of models of intermediate resolution aiming at balancing computational cost and physiological resolution. These models are cell-based and referred to as the Kirchhoff network model (KNM, [34]), and as the associated simplified Kirchhoff network model (SKNM, [35]). KNM and SKNM are based on representing individual cells and are similar to previous models of small collections of cardiomyocytes, see, e.g., [36, 37].

The aim of this paper is to evaluate the computational demands and physiological accuracy of the five models, EMI, BD, MD, KNM, and SKNM. Initially, we determine the mesh resolutions required for numerically accurate solutions. These resolutions form the basis for assessing the computational needs of each model. Subsequently, we analyze how these models simulate the electrochemical wave in the vicinity of an infarcted area. Notably, the models exhibit marked variations in conduction velocity as the excitation wave traverses the border zone between healthy and infarcted tissue. Such variations are critical, as they may lead to divergent interpretations about the presence of reentry waves, which in turn could be the precursors of life-threatening arrhythmias. Additionally, we present a case where the sub-cellular resolution of the EMI model is essential to demonstrate how a non-uniform distribution of certain ion channels along the cell membrane can significantly alter membrane dynamics. Lastly, we explore potential enhancements to our EMI code to increase its efficiency. While we have managed to reduce the computing time, the EMI model remains computationally intensive. Further development of more efficient solvers is imperative for using the EMI model in simulating large tissues effectively.

## 2 Methods

In this section, we briefly present the models, EMI, BD, MD, KNM and SKNM. We start by the EMI model and base the presentation of the other models on the EMI formulation.

### 2.1 The extracellular-membrane-intracellular model (EMI)

In the EMI model, the spatial domain consists of a number of cells, 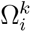, and a surrounding extracellular space, Ω_*e*_ (see Figure 1a). The interface between 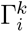 and Ω_*e*_ defines the cell membrane, Γ_*k*_. In addition, the interface between two neighboring cells, 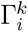 and 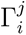, defines an intercalated disc, Γ_*k,j*_. The EMI model for the electrical potential in and surrounding the excitable cells can be expressed by the following system of equations (see, e.g., [12, 10]),

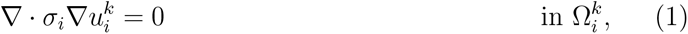

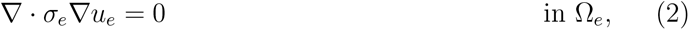

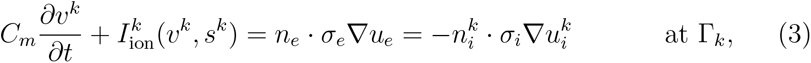

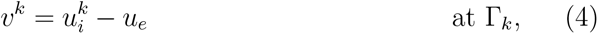

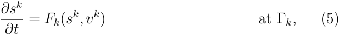

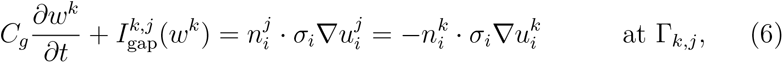

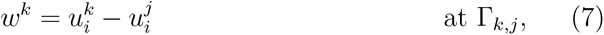

for each cell *k* with neighbors *j*. The unknown variables of the model are the intracellular potentials in each cell, 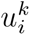 (in mV), defined in 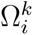, the extracellular potential surrounding the cells, *u*_*e*_ (in mV), defined in Ω_*e*_, the membrane potential of each cell, *v*^*k*^ (in mV), defined at the membrane, Γ_*k*_, and the intercalated disc potentials, *w*^*k*^, defined at the intercalated discs, Γ_*k,j*_.

**Figure 1:**
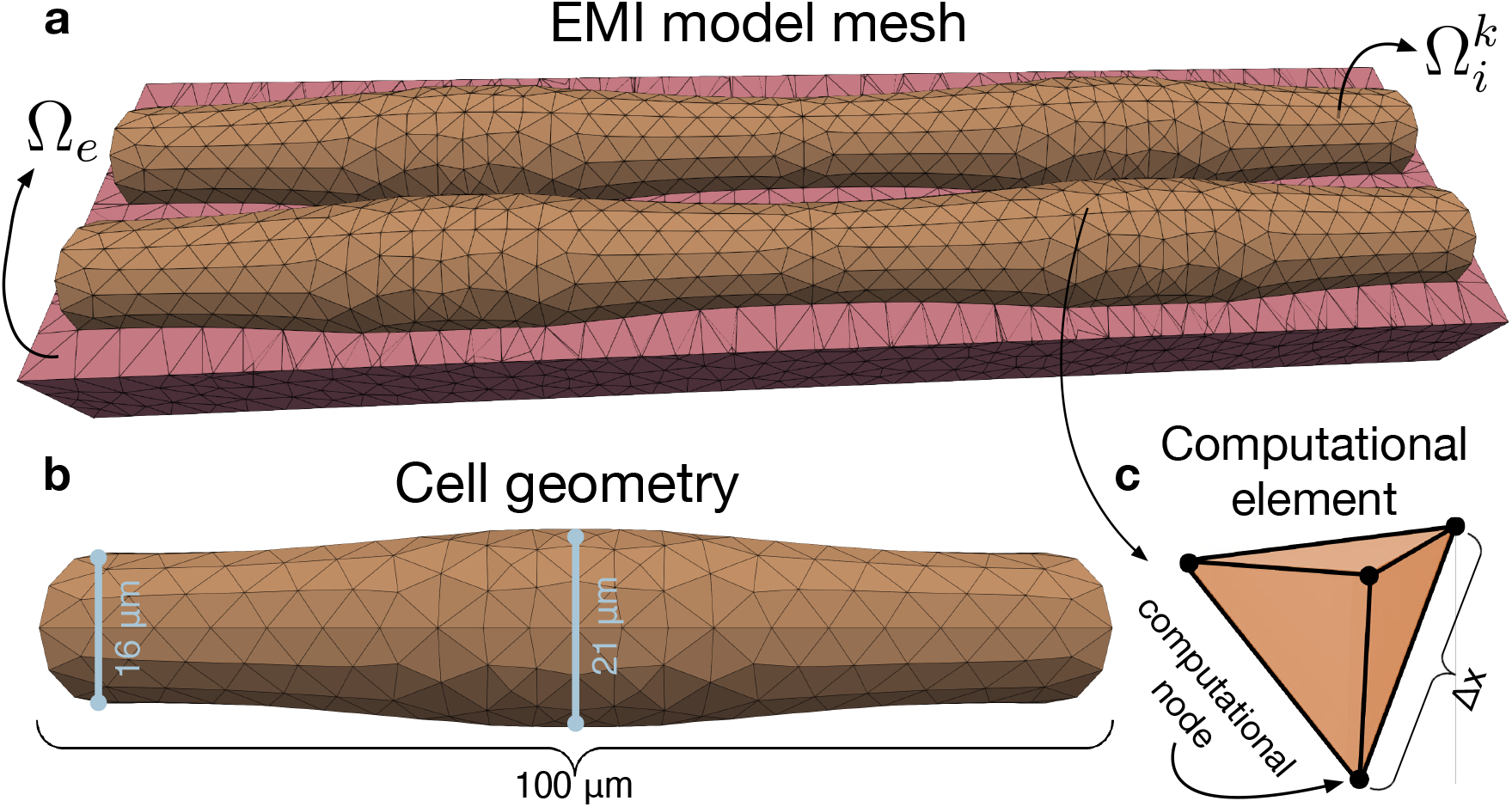
Illustration of the components of the computational mesh for the EMI model. **a**: Illustration of an EMI model mesh for 2*×*2 connected cells. The orange volumes are the cells, 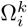, and the pink volume is the extracellular space, Ω_*e*_. The cell membrane, Γ_*k*_, is defined at the boundary surface between the intracellular and extracellular spaces, and the intercalated discs, Γ_*k,j*_, are defined at the boundary surface between two cell volumes, 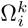 and 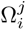. Note that the extracellular space surrounds the cells on all sides, but the upper half of the extracellular volume is removed from the illustration to make the cells visible. **b**: Illustration of the geometry of the cells. The cells are shaped as cylinders with length *l*_*x*_ = 100 *µ*m and diameter ranging between 16 *µ*m at the cell ends and 21 *µ*m at the center of the cells. **c**: Illustration of one computational element in the EMI model mesh. Each element is made up of four computational nodes with a typical distance of Δ*x*.

In addition, a number of additional state variables, *s*^*k*^, are defined at the membrane, representing ionic concentrations and gating variables for the ion channels used to compute the ionic membrane current density, 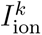 (in *µ*A/cm^2^). The dynamics of these state variables are modeled by *F*_*k*_. We use ion the adult ventricular base model from [38] for *F*_*k*_ and 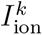 The formulation of this model is found in the Supplementary Information.

The parameters of the EMI model are the intracellular conductivity, *σ*_*i*_ (in mS/cm), the extracellular conductivity, *σ*_*e*_ (in mS/cm), the membrane capacitance, *C*_*m*_ (in *µ*F/cm^2^), and the intercalated disc capacitance, *C*_*g*_ (in *µ*F/cm^2^). Furthermore, the current density between neighboring cells, 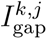, is modeled by a simple passive model

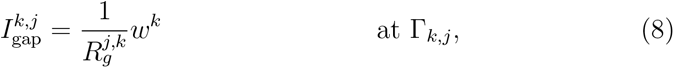

where 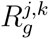 (in kΩcm^2^) is the resistance of the gap junctions connecting cells *k* and *j*.

The default parameter values used in our simulations are provided in Table 1. The cells are shaped as cylinders with length *l*_*x*_ = 100 *µ*m and radius ranging between 8 *µ*m at the cell ends and 10.5 *µ*m at the center of the cells (see Figure 1b). The distance between cell centers in the *y*-direction is *l*_*y*_ = 20 *µ*m. The minimum distance between the intracellular space to the outer boundary of the extracellular space is 5 *µ*m.

**Table 1:** Default parameter values used in the EMI model simulations. Here, *l*_*x*_ and *l*_*y*_ are the cell length in the *x*-direction and the cell width in the *y*-direction, respectively. In addition, *L*_*z*_ is the width of the entire domain (both intracellular and extracellular) in the *z*-direction.

| Parameter | Value | Parameter | Value |
| --- | --- | --- | --- |
| $C_m$ | $1 \mu\text{F}/\text{cm}^2$ | $R_g$ | $0.003 \text{ k}\Omega \text{ cm}^2$ |
| $C_g$ | $0.5 \mu\text{F}/\text{cm}^2$ | $l_x$ | $100 \mu\text{m}$ |
| $\sigma_i$ | $4 \text{ mS}/\text{cm}$ | $l_y$ | $20 \mu\text{m}$ |
| $\sigma_e$ | $20 \text{ mS}/\text{cm}$ | $L_z$ | $31 \mu\text{m}$ |

### 2.2 The bidomain model (BD)

In the bidomain model (BD), the tissue is not separated into distinct intracellular and extracellular parts as in the EMI model. Instead, both the intracellular space, the membrane and the extracellular space are assumed to exist everywhere in the domain (see Figure 2a). Based on this assumption, a formulation of BD can be derived from the EMI model (see [7]). The bidomain model reads

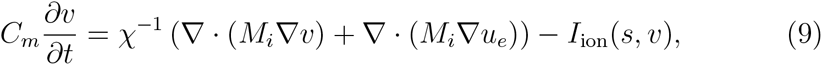

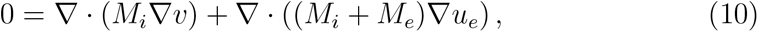

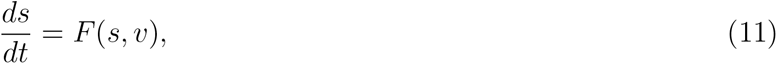

where *v* (in mV) is the membrane potential, and *u*_*e*_ (in mV) is the extracellular potential. In addition, *I*_ion_, *s* and *F* are the ionic current density, additional state variables and the dynamics of the additional state variables, modeled by [38] like for the EMI model. The BD parameters can be defined based on the EMI model mesh and parameters (see [7]). In particular, *χ* is the membrane surface to volume ratio, which can be computed by

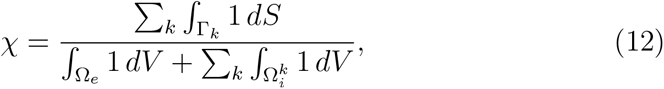

from the EMI model mesh. Furthermore, for a two-dimensional collection of cells

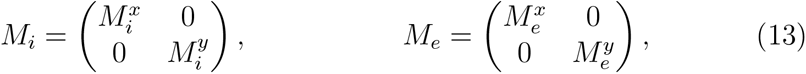

where

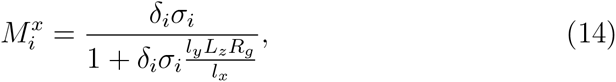

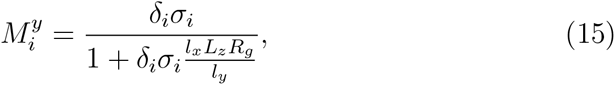

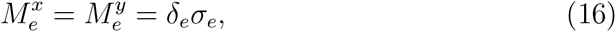

and *l*_*x*_ is the cell length, *l*_*y*_ is the cell width, *L*_*z*_ is the width of the domain in the *z*-direction, and *δ*_*i*_ and *δ*_*e*_ are the intracellular and extracellular volume fractions, respectively. These volume fractions can be computed from the EMI model mesh by

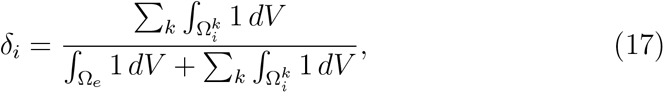

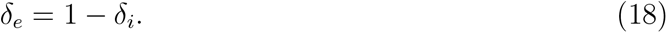

**Figure 2:**
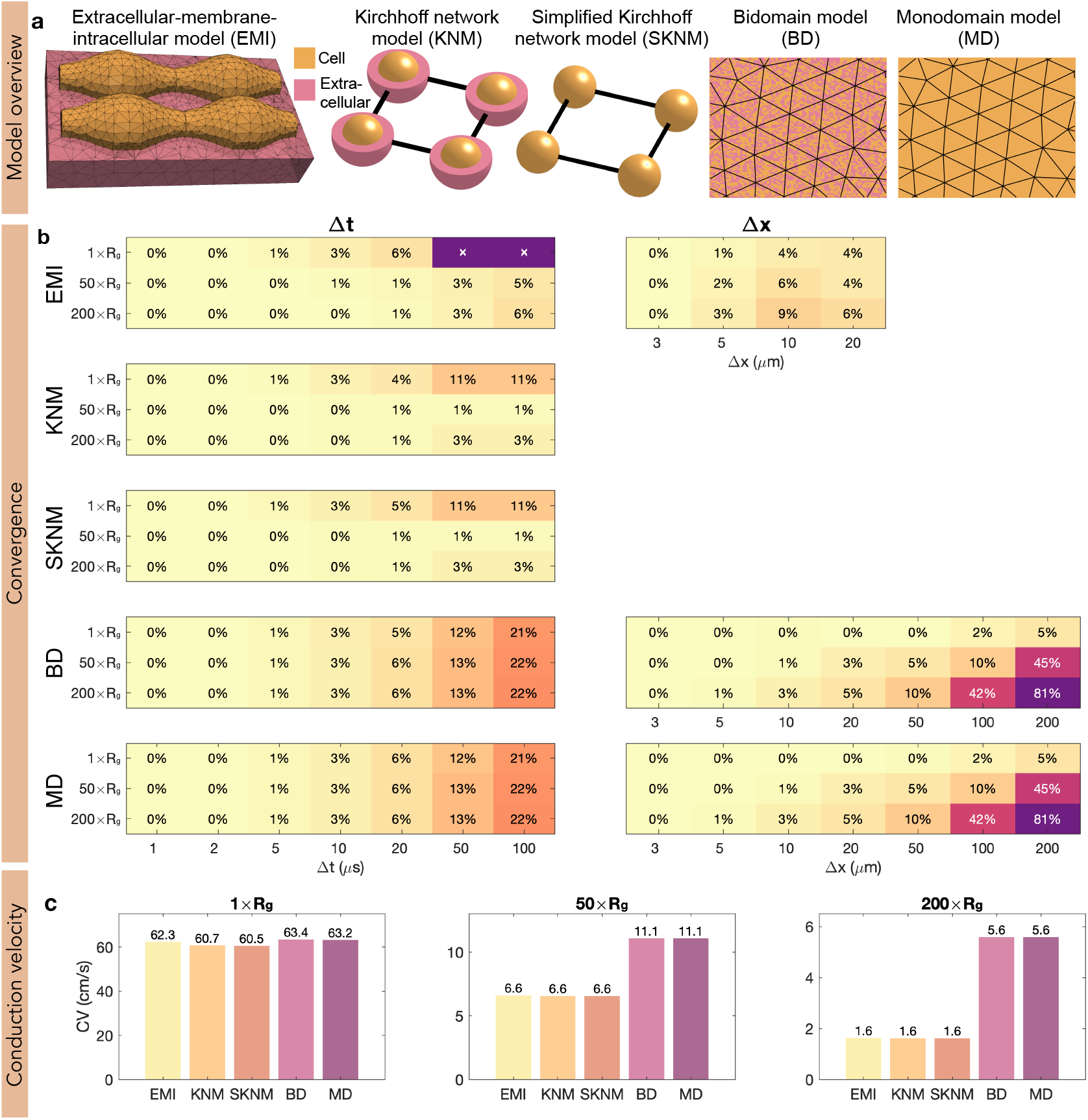
Model overview and convergence analysis. **a**: Overview of the five considered models of cardiac conduction: EMI, KNM, SKNM, BD, and MD. In EMI, the extracellular space, the membrane and the cells are spatially resolved in 3D. In KNM, each cell and associated extracellular space are represented as single node points, and in SKNM, the representation is simplified to only represent each cell. In BD, the tissue is treated as a continuum consisting of both extracellular space, intracellular space and membrane everywhere. In MD, the BD representation is simplified to only represent the cell membrane. **b**: Convergence analysis for the five models. We compute the conduction velocity (CV) along a strand of 15 cells for each model for a number of different spatial (Δ*x*) and temporal (Δ*t*) resolutions and report the difference (in percent) from the CV computed using the finest considered resolution (Δ*t* = 1 *µ*s, Δ*x* = 3 *µ*m). We consider three different values of the gap junction resistance, the default value, 1 *× R*_*g*_, and 50 *× R*_*g*_ and 200 *× R*_*g*_. Note that there is no spatial resolution parameter for KNM and SKNM since the position of the cells define the node points. Furthermore, the operator splitting algorithm [23] used to solve the EMI model is unstable the default value of *R*_*g*_ (1 *× R*_*g*_) and Δ*t ≥* 50 *µ*s. When adjusting one of the discretization parameters, the other is fixed at the finest resolution. **c**: Conduction velocity computed using the finest resolution of each model for three different values of the gap junction resistance, *R*_*g*_.

For a one-dimensional collection of cells, we have

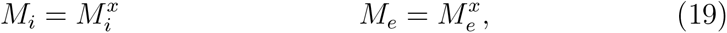

where 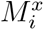 and 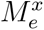 are given by (14) and (16) with the cell width *l*_*y*_ replaced by the domain width *L*_*y*_, which is equal to the cell width plus the width of the extracellular space (30 *µ*m in our 1D simulations).

### 2.3 The monodomain model (MD)

The monodomain model (MD) can be derived from the BD model based on the assumption

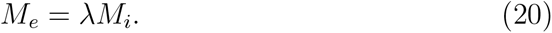

The model then reads

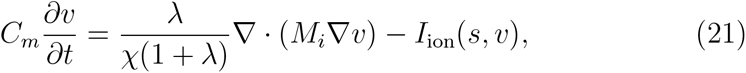

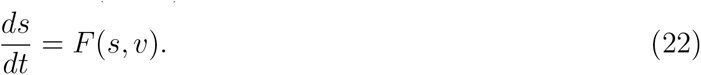

In our 1D strand simulations where (20) holds, we use the definition

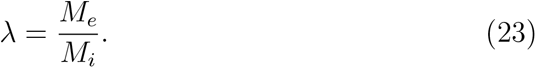

In the 2D simulations where the assumption (20) do not hold, we use the approximation of *λ* from [35], i.e.,

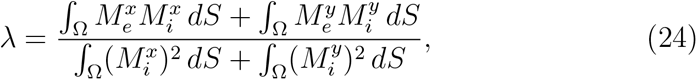

where Ω is the computational domain and 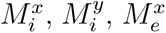, and 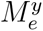 are defined in (14)–(16).

### 2.4 The Kirchhoff network model (KNM)

In the Kirchhoff network model (KNM), each cell in the excitable tissue is represented by a single nodal point, as opposed to being spatially resolved as in the EMI model (see Figure 2a). In addition, the extracellular space surrounding each cell is represented in each node. The model is derived by applying Kirchhoff’s current law for each nodal point [34] and reads

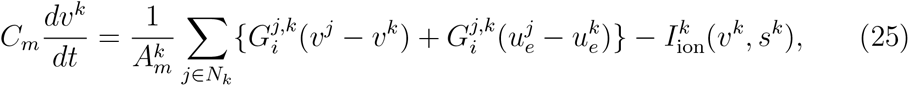

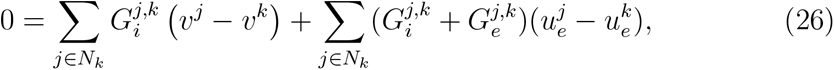

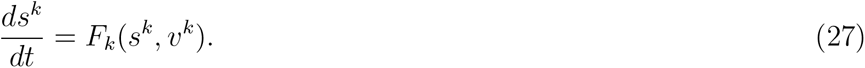

for all *k* numbering the cells in the tissue. Here, *v*^*k*^ is the membrane potential *e* of cell *k* (in mV), 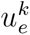 is the associated extracellular potential (in mV), and *N*_*k*_ is a collection of all the neighboring cells of cell *k*. Furthermore, 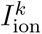, *s*^*k*^ and *F*_*k*_ are the ionic current density, additional state variables and the dynamics of the additional state variables, modeled by [38] like for the EMI model. Moreover, 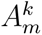 (in cm^2^) is the membrane area of cell *k*, and can be computed from the EMI model mesh by

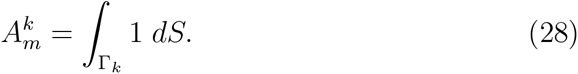

The parameters 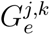 and 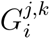 are conductances defined by,

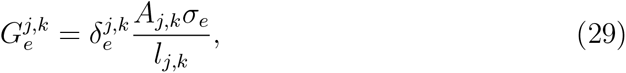

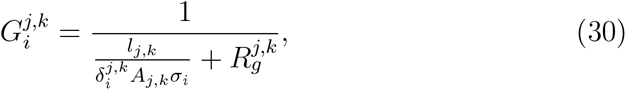

where 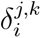 and 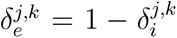 are the extracellular and intracellular volume fractions, respectively, associated with cell *k*. These are assumed to be constant throughout the domain and defined by (17) and (18). In addition, *A*_*j,k*_ is the cross-sectional area (both intracellular and extracellular) between the centers of cell *k* and *j* and *l*_*j,k*_ is the distance between the centers. If the cells are connected in the *x*-direction, then *A*_*j,k*_ = *l*_*y*_*L*_*z*_ (or *A*_*j,k*_ = *L*_*y*_*L*_*z*_ in the 1D case) and *l*_*j,k*_ = *l*_*x*_. Similarly, if the cells are connected in the *y*-direction, then *A*_*j,k*_ = *l*_*x*_*L*_*z*_ and *l*_*j,k*_ = *l*_*y*_.

### 2.5 The Simplified Kirchhoff network model (SKNM)

The Simplified Kirchhoff network model (SKNM) can be derived from KNM based on an assumption similar to that used to define MD from BD [35]. More specifically, based on the assumption

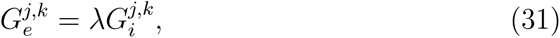

KNM simplifies to

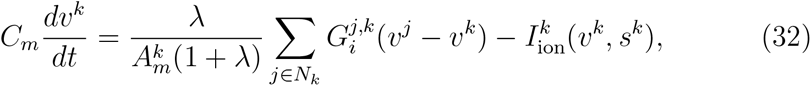

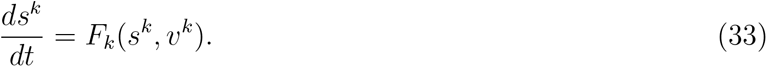

In our 1D strand simulations where (31) holds, we use the definition

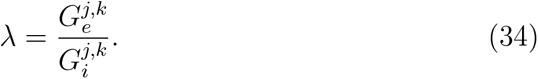

In the 2D simulations where the assumption (31) do not hold, we use the approximation of *λ* from [35], i.e.,

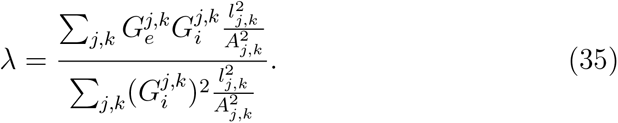

### 2.6 Implementation

In this subsection we describe the implementation of the numerical solution algorithms for the five models EMI, KNM, SKNM, BD, and MD.

#### 2.6.1 Implementation of KNM and SKNM

The numerical solution of KNM and SKNM are implemented in C++. A classical first-order operator splitting scheme is used to split the linear and the non-linear parts of the systems (see, e.g., [39]). Here the non-linear part of the systems is the part concerning the membrane dynamics, 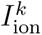 and *F*_*k*_. These non-linear ordinary differential equations are solved using a first-order Rush-Larsen method [40] with code generated using the Gotran code generator [41] and OpenMP parallelization [42]. The remaining linear equations are solved using the backward Euler method. The resulting linear systems are solved using the biconjugate gradient stabilized method (BiCGSTAB) from the MFEM finite element library [43, 44]. For KNM, we use a block Jacobi preconditioner [45]. Unless otherwise stated, we use the time step Δ*t* = 10 *µ*s in the solution of KNM and SKNM.

#### 2.6.2 Implementation of BD and MD

The numerical solution of BD and MD are also implemented in C++ using the MFEM finite element library [43, 44]. The meshes are generated using gmsh [46], and we use first-order finite elements. Like for KNM and SKNM, we split the system of equations into linear and non-linear parts using a first-order operator splitting scheme [39]. Again, the non-linear part of the systems is the part concerning the membrane dynamics, *I*_ion_ and *F*, and these are solved using a first-order Rush-Larsen method [40] with code generated using the Gotran code generator [41] and OpenMP parallelization [42]. The remaining linear equations are solved using the backward Euler method and the linear systems are solved using the generalized minimal residual method (GMRES). For BD, we use a block Jacobi preconditioner [45]. Unless otherwise stated, we use the time step Δ*t* = 10 *µ*s and the spatial resolution Δ*x* = 10 *µ*m in the solution of BD and MD.

#### 2.6.3 Implementation of EMI

The EMI meshes are generated using gmsh [46]. To solve the EMI equations, we apply the temporal and spatial splitting algorithm introduced in [23] (see Algorithm 1). In this splitting algorithm, the non-linear membrane dynamics (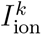 and *F*_*k*_) are treated in Step 1 like in the operator splitting for KNM, SKNM, BD and MD. We use a first-order Rush-Larsen method [40] with code generated using the Gotran code generator [41] and OpenMP parallelization [42] to solve these ordinary differential equations.

In addition, operator splitting techniques are applied so that in each time step, the linear system of model can be split spatially into distinct parts that can be handled separately [23]. More specifically, the linear system is split into one equation for each cell (Step 2) that may be treated independently of each other (i.e., in parallel). In addition, we get one system for the extracellular part of the domain that can be handled separately (Step 3). Consequently, the sub-problems needed to be solved for each time step are standard Laplace problems with Dirichlet, Neumann or Robin boundary conditions. We have used one iteration of this algorithm for each time step, but the accuracy of the splitting algorithm could potentially be improved by increasing the number of iterations (see [23]).

##### Algorithm 1

Spatial and temporal splitting algorithm applied to solve the EMI model equations (1)–(7), taken from [23, 47]. Here, 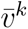, *ū*_*e*_, 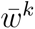, and 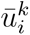 are temporary solutions of *v*^*k*^, *u*_*e*_, *w*^*k*^, and 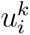 respectively. Moreover, *N*_it_ and *M*_it_ denote the number of outer and inner iterations of the algorithm. In our computations, we use *N*_it_ = *M*_it_ = 1.

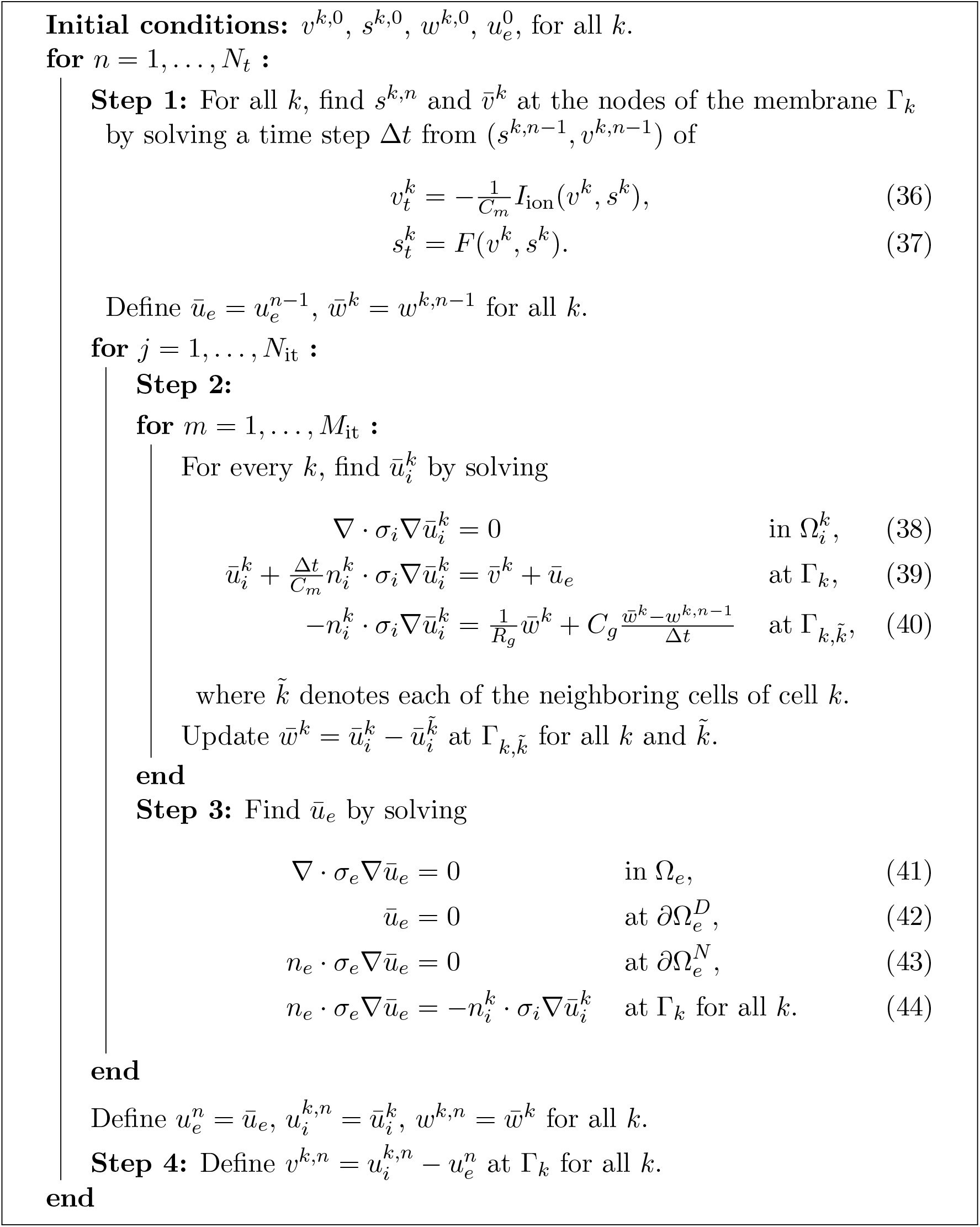

In our implementation, we use first-order finite elements from the MFEM finite element library [43, 44]. The resulting linear systems for the intra-cellular and extracellular systems are solved using the conjugate gradient method (CG) with a Gauss-Seidel preconditioner [45]. We also consider the use of CG with an algebraic multigrid (AMG) preconditioner, namely the BoomerAMG implementation from *hypre* [48], as well as two different sparse direct solvers, UMFPACK [49] and SuperLU [50]. BoomerAMG and SuperLU are parallelised using OpenMP, whereas the other solvers do not benefit from multithreading. The intracellular systems are solved in parallel using OpenMP [42]. Unless otherwise stated, we use the time step Δ*t* = 10 *µ*s and the spatial resolution Δ*x* = 5 *µ*m in the solution of EMI.

## 3 Results

### 3.1 Convergence of model solutions

In Figure 2b, we investigate the spatial and temporal resolutions required to obtain converged numerical solutions for the five models EMI, KNM, SKNM, BD, and MD. For each considered resolution and model, we compute the conduction velocity (CV), compare it to the CV computed for the considered model using the finest resolution (Δ*t* = 1 *µ*s, Δ*x* = 3 *µ*m), and report the difference in percent. Since we will use the models to study an example of an infarction scar with a border zone of reduced gap junction coupling (increased *R*_*g*_), we investigate convergence both for the default gap junction resistance, 1 *× R*_*g*_, and for two cases of increased resistance, 50 *× R*_*g*_ and 200 *× R*_*g*_, similar to what was done in [34].

#### 3.1.1 Temporal resolution

In the left panel of Figure 2b, we investigate the effect of the time step, Δ*t*, used in the numerical solution of the models. We observe that in order to get an error of 3% or lower in all cases, we need a time step of Δ*t* = 10 *µ*s or smaller. Note that for the EMI model, the operator splitting algorithm used to solve the system of equations [23, 47] is unstable when the two largest considered time steps are used for the default value of *R*_*g*_.

#### 3.1.2 Spatial resolution

In the right panel of Figure 2b, we investigate the effect of the spatial resolution, Δ*x*, used in the model meshes. Note here that for KNM and SKNM, there is no tuneable spatial discretization parameter since the location of the cells dictates the node points. Note also that for EMI, a relatively fine mesh (Δ*x* = 20 *µ*m) is needed merely to represent the geometry of the cells and the surrounding extracellular space, so no larger values of Δ*x* are considered. Furthermore, to achieve an error smaller than or equal to 3%, a 5 *µ*m resolution seems to be required for the EMI model. For BD and MD, the errors associated with choosing a large value of Δ*x* seems to be most pronounced for increased values of *R*_*g*_. Moreover, in order to get make the error smaller than 3% for all the considered values of *R*_*g*_, a resolution of 10 *µ*m appears to be needed for BD and MD.

In the remaining simulations in this study, we use Δ*x* = 5 *µ*m for EMI, Δ*x* = 10 *µ*m for BD and MD, and Δ*t* = 10 *µ*s for all models.

### 3.2 Comparison of model solutions for EMI, KNM, SKNM, BD and MD

In Figure 2c, the CV computed using the finest resolution of all the considered models (EMI, KNM, SKNM, BD, and MD) are reported for three values of the gap junction resistance, *R*_*g*_. Here we observe that the CVs computed using all five models are quite similar for the default value of *R*_*g*_. However, when *R*_*g*_ is increased, EMI, KNM and SKNM appears to be very similar, and BD and MD are very similar, but there are some differences in the CV between these two groups of models, consistent with what was found in [7, 34].

#### 3.2.1 Infarcted region

In Figure 3, we further investigate the difference between the models using an example application of a small infarction scar. Figure 3c shows the set-up used in the simulation. We consider a collection of 13*×*65 cells, and each of the small rectangles in Figure 3c represents a cell in the tissue. The volume of each cell is about 30 pL, see Figure 1. The pink and yellow areas both represent healthy tissue, and a stimulation current is applied in the yellow area. The purple area represents the scar core. Here, the gap junction resistance, *R*_*g*_, is set to infinity and the conductance of the sodium channels, *g*_Na_ is set to zero, representing injured, non-conductive tissue. The blue area surrounding the scar represents the scar border zone. In this area, the extracellular potassium concentration and the gap junction resistance are increased compared to the healthy case. More specifically, *R*_*g*_ is increased by a factor linearly increasing from 50 at the outer part of the border zone to 200 at the inner part. Similarly, the extracellular potassium concentration increases linearly from 5.4 mM to 10 mM from the outer to the inner parts of the border zone.

**Figure 3:**
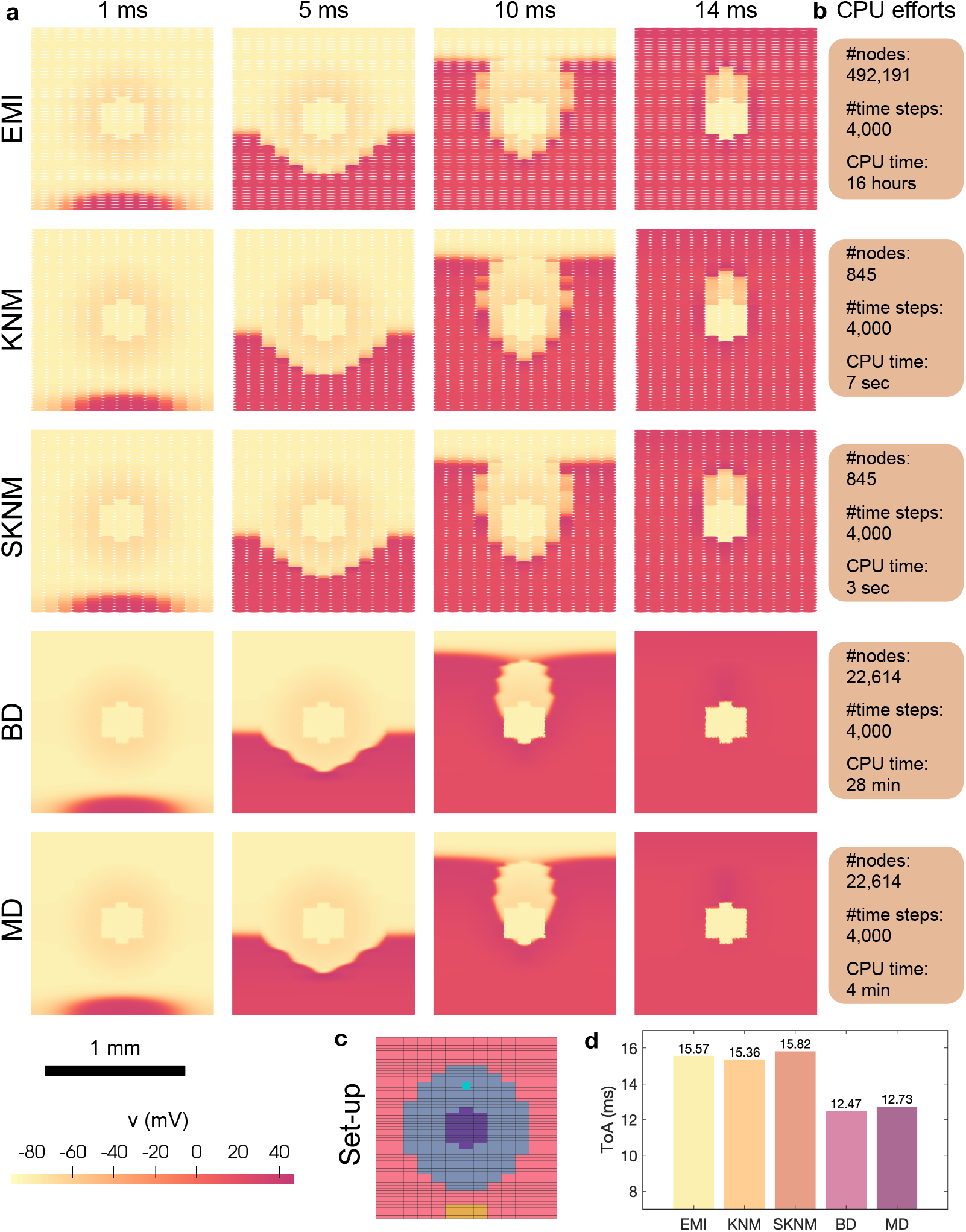
Simulations of a small domain with an infarction scar and border zone. **a**: Snapshots of the solutions at four points in time after a stimulation is applied for EMI, KNM, SKNM, BD and MD. **b**: CPU efforts of the simulations in terms of number of nodes, number of time steps and CPU time required to run a 40 ms simulation. **c**: Applied domain set-up. Each cell is represented by a small colored rectangle. The pink area represents healthy tissue, the blue area represents infarction border zone and the purple area represents the scar core. The yellow area marks the stimulation site (in the healthy region), and the turquoise point marks a measurement point used in Panel d. The tissue consists of 13*×*65 cells and is 1.3 mm*×* 1.3 mm. **d**: Point in time when the membrane potential in the turquoise point in Panel c reaches *v ≥* 0 mV for each of the five models. We use Δ*x* = 3 *µ*m for EMI, Δ*x* = 10 *µ*m for BD and MD, and Δ*t* = 10 *µ*s for all models.

In Figure 3a, we show snapshots of the solution of all the five models (EMI, KNM, SKNM, BD, and MD) at four points in time after stimulation is applied for a simulation using this set-up. When the excitation wave travels in the healthy part of the tissue (two leftmost panels), all five models appears to provide very similar solutions. However, as the wave reaches the high-resistance border zone (rightmost two panels), the wave clearly travels faster for BD and MD than for EMI, KNM and SKNM. Moreover, Figure 3d shows the point in time after the stimulation was applied when the membrane potential in the turquoise point in Panel c first reaches a membrane potential above 0 mV. We observe that this time of arrival (ToA) is quite similar for EMI, KNM and SKNM, and similar for BD and MD, but that the ToA differs between the two groups of models.

### 3.3 Comparison of CPU times for EMI, KNM, SKNM, BD and MD

In Figure 3b, we report the number of mesh nodes, number of time steps and total CPU time for a 40 ms simulation using this set-up for each of the five models. We observe that the CPU time for SKNM (3 sec) is a bit shorter than the CPU time for KNM (7 sec) and considerably shorter than the CPU time for EMI (16 hours). Since the solution of these three models appears to be very similar, we will use SKNM as a representative model for EMI and KNM. Similarly, the CPU time for MD (4 min) is quite a bit shorter than that of BD (28 min), and we will therefore use MD as a representative model of MD and BD in simulations of a larger domain.

### 3.4 Comparison of SKNM and MD solutions around an infarction scar

To illustrate that the differences in conduction velocity (Figure 2c) and time of arrival (Figure 2d) between the cell-based models (EMI, KNM, SKNM) and the homogenized models (BD, MD) can significantly influence the simulation results, we consider an example of a larger infarction scar and compare the solution of SKNM and MD. The set-up is similar to that illustrated in Figure 3c, except that the radius of the scar core is increased to 2 mm and the width of the surrounding border zone is set to 1.5 mm. The total domain consists of 100*×*500 cells and is 1 cm *×* 1 cm. The scar core and border zone are treated in the same manner as in the simulations reported in Figure 3, except that the gap junction resistance increases by a factor from 20 to 200 instead of from 50 to 200 from the outer to the inner parts of the border zone. In order to investigate the tissue’s susceptibility to reentry, we apply a classical S1-S2 stimulation protocol (see, e.g., [51, 52]). The S1 stimulation is applied in 3*×*5 cells in the lower center of the domain like illustrated in Figure 3c, and the S2 stimulation is applied 310 ms later in the entire lower left quarter of the domain (except for in the scar core).

#### 3.4.1 Reentrant wave generated for SKNM, not for MD

In Figure 4, we show snapshots of the solution of SKNM and MD at five points in time after the S2 stimulation was applied in an infarction scar simulation using this set-up. We observe that for SKNM, a wave is generated traveling around the scar core in the counter-clockwise direction. Directly following the S2 stimulation, the wave is not able to move in the clockwise direction because the cells above the stimulation site are not yet depolarized.

**Figure 4:**
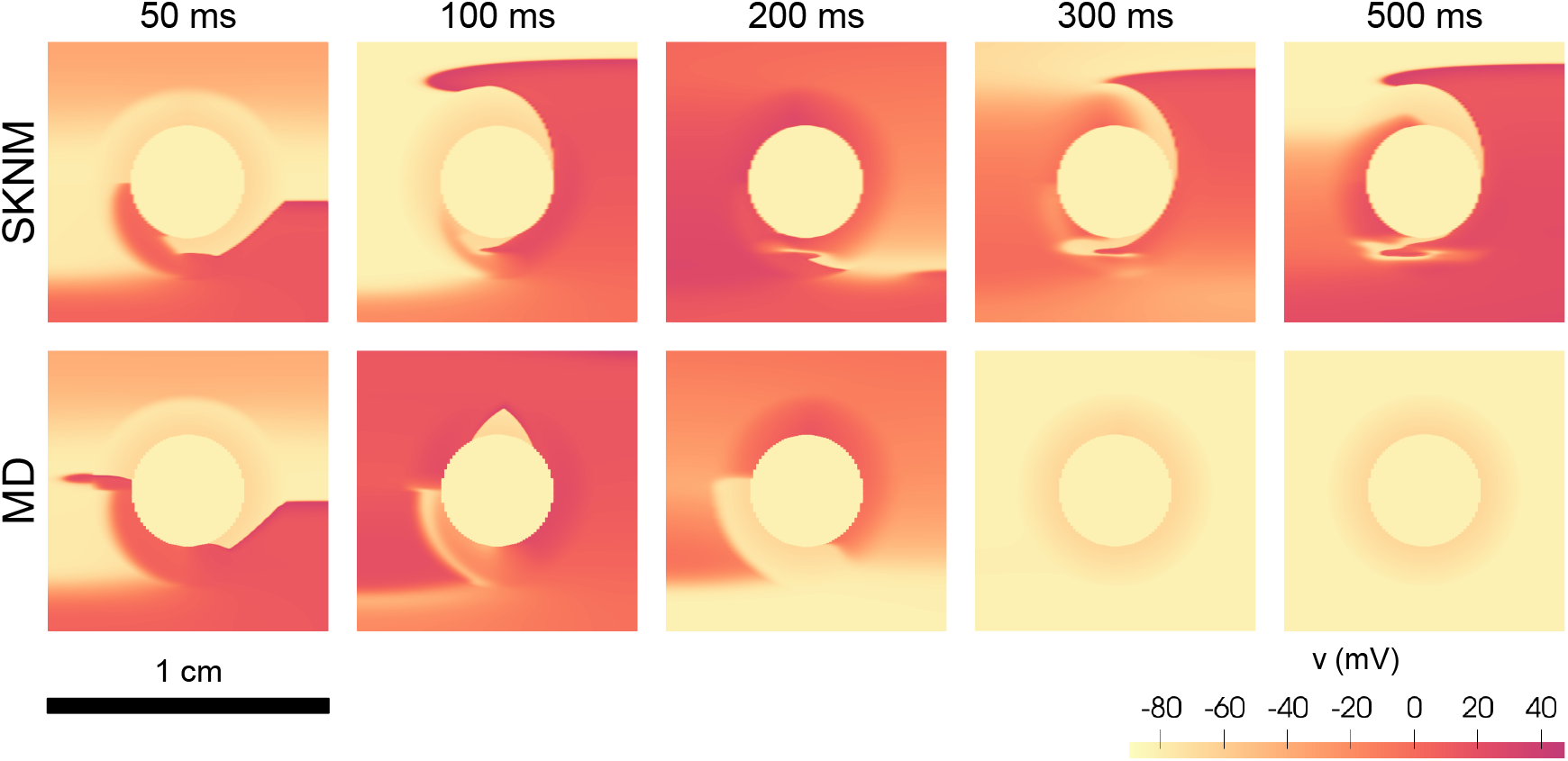
Reentrant wave generated around an infarction scar for SKNM, but not for MD. Reentry is attempted to be generated by applying an S1-S2 stimulation protocol [51, 52]. The S1 stimulation is applied in the lower part of the domain (see Figure 3c), and the S2 stimulation is applied in the lower left quarter of the domain (except for in the scar core) 310 ms after the S1 stimulation was applied. The time points at the top of the plots report the time after the S2 stimulation was applied. In the border zone, the extracellular potassium concentration increases linearly from 5.4 mM at the outer part to 10 mM in the inner part. Furthermore, *R*_*g*_ is increased by a factor that similarly increases linearly from 20 to 200. We use Δ*x* = 10 *µ*m for MD and Δ*t* = 10 *µ*s for both models.

When the wave reaches the area of the S2 stimulation after traveling one round, however, the tissue is repolarized enough for the wave to continue traveling around the scar core and we get a reentrant wave around the scar core.

For MD, on the other hand, the S2 stimulation induces a wave traveling both in the clockwise and counter-clockwise directions. This may be explained by the fact that the CV is faster for MD than for SKNM (Figure 2c and Figure 3a), and that the excitation wave following the S1 stimulation has traveled so fast that the tissue above the S2 stimulation area is able to depolarize again following the S2 stimulation. Since the wave travels in two directions, there is no tissue ready to be depolarized again after the wave has traveled through the entire border zone and healthy tissue, and, consequently, no reentrant wave is generated for MD.

#### 3.4.2 Reentrant spiral wave in the border zone for SKNM and MD

In Figure 4, we observed that a reentrant wave was generated around the scar core for SKNM, but not for MD. However, using another choice of parameters, a reentrant spiral wave is generated in the border zone for both models (see Figures 5 and 6). In this case, we let the gap junction resistance be increased by a factor ranging from 20 to 375 in the border zone instead of from 20 to 200 (linear increase from outer to inner parts of the border zone).

**Figure 5:**
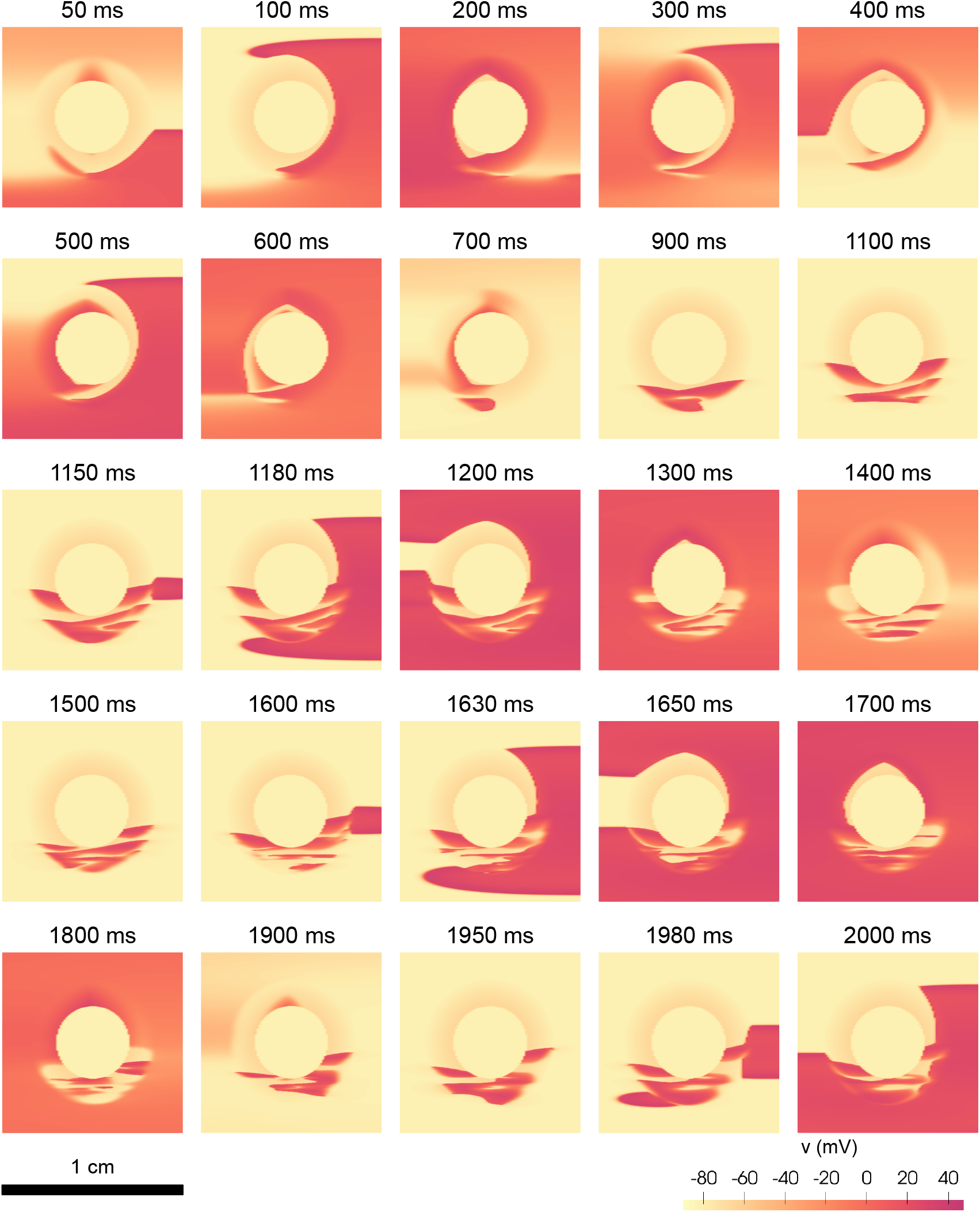
SKNM simulation of a reentrant spiral wave in the border zone. We apply an S1-S2 stimulation protocol [51, 52]. The S1 stimulation is applied in the lower part of the domain (see Figure 3c), and the S2 stimulation is applied in the lower left quarter of the domain (except for in the scar core) 310 ms after the S1 stimulation was applied. The time points at the top of the plots report the time after the S2 stimulation was applied. In the border zone, the extracellular potassium concentration increases linearly from 5.4 mM at the outer part to 10 mM in the inner part. Furthermore, *R*_*g*_ is increased by a factor that similarly increases linearly from 20 to 375. We use Δ*t* = 10 *µ*s.

**Figure 6:**
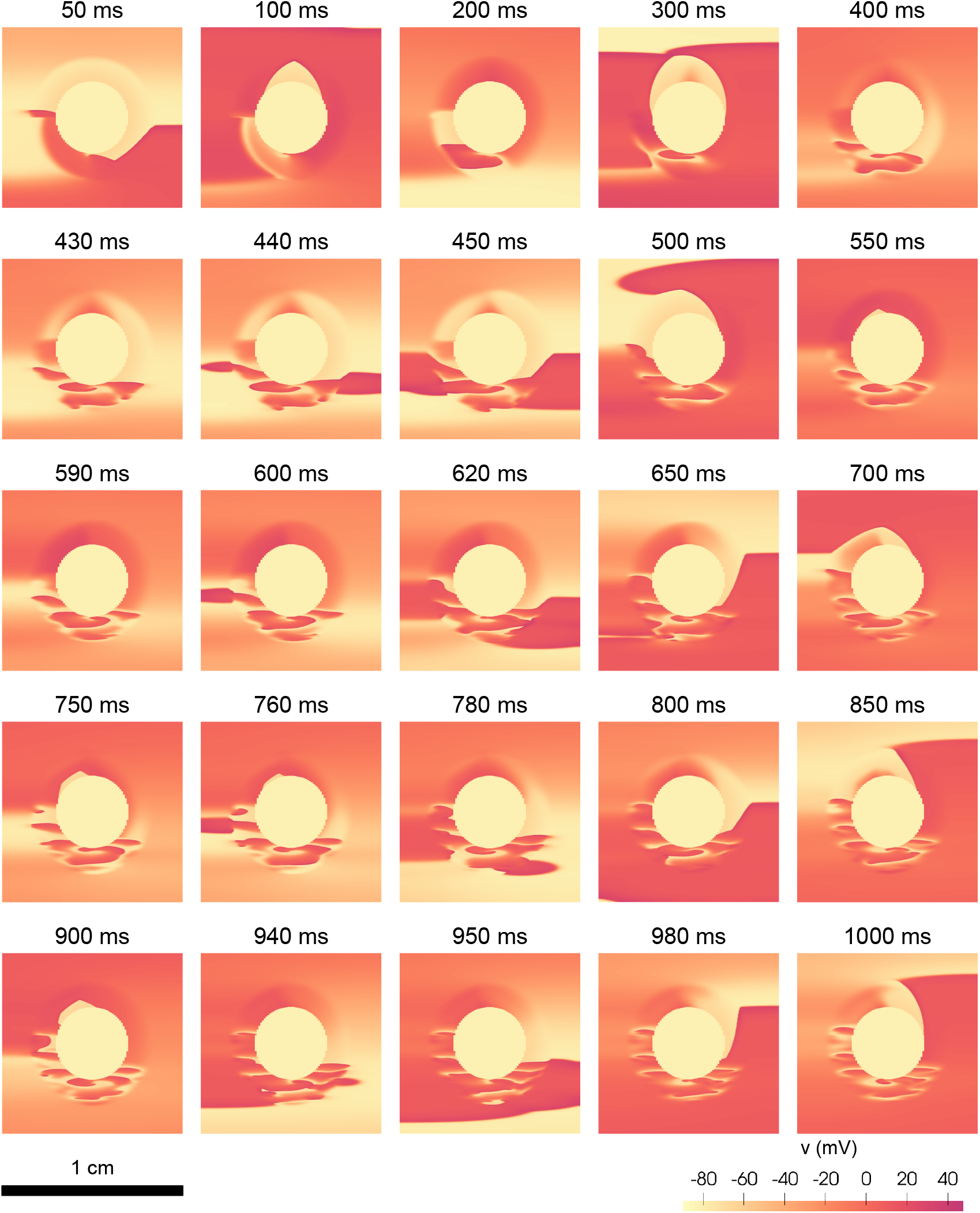
MD simulation of a reentrant spiral wave in the border zone. The simulation set-up is the same as that used in Figure 5. We use Δ*x* = 10 *µ*m and Δ*t* = 10 *µ*s.

In Figure 5, we show the activity following the S2 stimulation in the SKNM simulation. We observe that for the first 600 ms, a reentrant wave similar to that observed in Figure 4 is generated traveling around the scar core. After this, the reentrant wave in the healthy tissue seems to dissipate, while a small reentrant spiral wave continues in the border zone. After some time (at 1150 ms after the S2 stimulation), this electrical activity in the border zone spreads to the healthy tissue and depolarizes it. Interestingly, the activity in the border zone continues even after the cells in the healthy tissue repolarize again and after some time (at 1600 ms and at 1980 ms) new waves spread from the border zone to the healthy tissue.

In Figure 6, the same simulation is conducted using MD. We observe that even though there are significant differences between the SKNM and MD solutions (Figure 5 and Figure 6), there are also qualitative similarities between the solutions. For example, Figure 6 shows that in this case, like for SKNM, a small reentrant spiral wave is generated in the border zone also in the MD solution. Furthermore, like in the SKNM simulation, excitation waves are repeatedly emitted from the border zone to the healthy tissue.

### 3.5 Modeling capability exclusive to EMI — effect of a non-uniform distribution of ion channels on the cell membrane

In Figures 2 and 3, we observed that KNM and SKNM provided good approximations of the EMI solution. Nevertheless, in some cases, the improved complexity of EMI makes the model able to accurately represent properties that cannot be represented in the more simple KNM and SKNM. One example of such an application is illustrated in Figure 7. In this case, we consider the case of a non-uniform distribution of ion channels along the cell membrane. More specifically, we compare the case when all the L-type calcium channels are uniformly distributed throughout the membrane (U), to the non-uniform (NU) case where the channels are located in five bands along the length of the cell (see Figure 7a). Figure 7b shows the action potentials computed in single-cell EMI model simulations using these two channel distributions, and we observe that the duration of the action potential is considerably reduced in the NU case. Such investigations would not be possible to conduct using KNM and SKNM, since the cells are not spatially resolved in those models.

**Figure 7:**
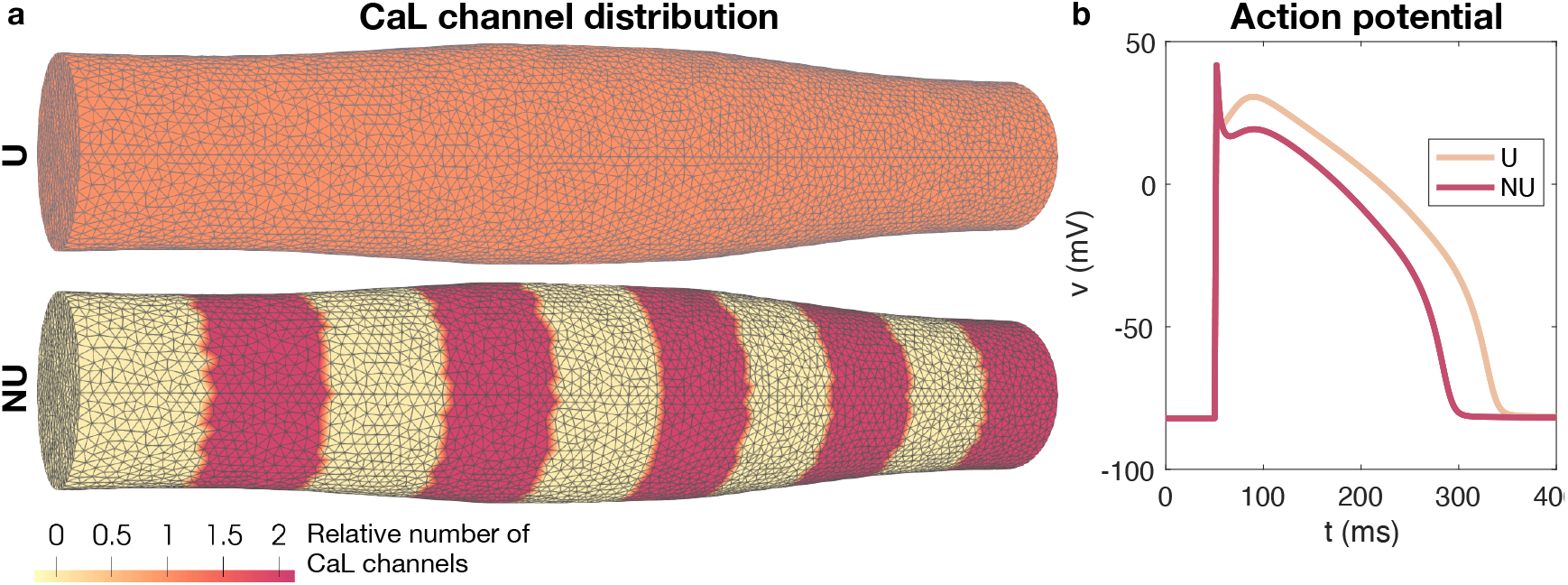
Comparison of a uniform (U) and non-uniform (NU) distribution of L-type calcium channels in a single-cell EMI model simulation. **a**: Illustration of the uniform and non-uniform distributions of L-type calcium channels in the EMI model membrane mesh. **b**: Action potentials computed using the EMI model for the two channel distributions, U and NU. We use Δ*x* = 1 *µ*m and Δ*t* = 0.01 ms. The membrane potential is plotted for three membrane points (one at each cell end and one in the center of the cell) for each of the channel distributions, but the solutions for the different membrane points are indistinguishable.

### 3.6 Parallel solvers for EMI

As noticed in Figure 3 above, the computation time for the EMI solver is very significantly longer than for the other models. This is reasonable since EMI offers sub-cellular resolution whereas the other models average many cells (MD, BD) or represent every cell (KNM, SKNM) without sub-cellular resolutions. Since EMI clearly is the most accurate representation of the biophysics involved, it is of great importance to speed up the simulations. In Algorithm 1, the EMI model has been broken down to components that invite parallel solution in the sense that all computations can be performed independently for every cell except for the extracellular potential that binds everything together. In this section, we consider steps to speed up this part of the computations. In particular, we describe numerical experiments concerned with the shared-memory parallel performance of the EMI solver implementation. The experiments are carried out on a 24-core AMD Epyc 7413 CPU.

Recall that the problem described in Section 3.3 requires about 16 hours to simulate a configuration of 13*×*65 cells for 4 000 time steps, using a sequential CG-Gauss-Seidel solver for the linear system arising from the extracellular equation each time. A profiling shows that more than 80 % of the execution time is spent on solving the extracellular equations (41)–(44), about 10 % on solving the intracellular equations (38)–(40), a mere 1 % on solving the membrane model (36)–(37), and a small, remaining fraction is mostly spent on moving data and updating the solution between time steps and solver iterations.

Based on this observation, we first consider alternative solvers for the extracellular equations. In addition to CG with Gauss-Seidel preconditioning, we also consider hypre (i.e., CG with AMG preconditioning) as well as the sparse direct solvers UMFPACK and SuperLU. Table 2 shows execution times for solving the extracellular equations for 10 time steps using the different solvers. Only sequential execution times are reported for CG-Gauss-Seidel and UMFPACK, whereas parallel execution times using up to 24 threads are reported for hypre and SuperLU.

**Table 2:** Execution time (in seconds) for solving the extracellular equations. We perform 10 time steps of the EMI model using the same setup as in Figure 3, where the extracellular system consists of 453 572 unknowns. Four different solvers are used, and we report the total execution time for all time steps. The measurements are obtained on a 24-core AMD Epyc 7413 CPU, where up to 24 threads are used if the solver is parallel (i.e., CG-AMG and SuperLU). Otherwise, only the serial execution time is reported.

| Threads | CG-Gauss-Seidel | CG-AMG (hypre) | UMFPACK | SuperLU |
| --- | --- | --- | --- | --- |
| 1 | 127.90 | 21.26 | 15.69 | 25.10 |
| 2 | – | 12.34 | – | 18.77 |
| 4 | – | 7.60 | – | 15.71 |
| 8 | – | 6.39 | – | 15.82 |
| 16 | – | 5.04 | – | 16.43 |
| 24 | – | 5.23 | – | 17.37 |

#### Speed-up using multiple threads

In the sequential case, hypre, UMFPACK and SuperLU yield speedup factors of 6.0, 8.2 and 5.1, respectively, over the baseline solver with CG and Gauss-Seidel preconditioning. SuperLU benefits from using up to 4 or 8 threads, which results in about 60 % improvement over its sequential runtime. At this point, SuperLU reaches the same performance as UMFPACK, about 15.7 seconds. Additional threads do not provide any further benefit and only result in poor parallel efficiency. By comparison, hypre benefits from using up to 16 threads and attains a speedup of 4.2*×* over its sequential runtime. It is thus the fastest among the candidates considered for solving the extracellular equations in parallel, and it results in a speedup of 25*×* compared to the sequential baseline solver.

#### Direct vs. iterative solvers

In the case of UMFPACK and SuperLU, the reported times include both the initial factorization as well as solving triangular linear systems during each time step. Although factorization is performed only once, it accounts for nearly half of the runtime for UMFPACK in the current example. On the other hand, if the number of time steps increases, more time will be spent on performing the triangular solves, and the factorization time eventually becomes negligible. By this reasoning, direct solvers can be regarded as more competitive than they would seem at first. Supposing that we ignore factorization, then the execution time for UMFPACK is 8.5 seconds. So, even in this case, hypre remains the fastest alternative.

#### Parallel performance of EMI model

Besides the extracellular equations (41)–(44), parallel processing is also beneficial when solving the intracellular equations (38)–(40) and the membrane model (36)–(37). Table 3 shows the execution time when using up to 24 threads to solve the EMI model for 10 time steps. The top half of the table shows the initial parallel performance of the code, whereas the bottom half shows the parallel performance after applying certain code optimizations that are discussed in detail below. The execution times are also broken down into five main parts, including solving the membrane model, as well as assembling and solving linear systems for the intracellular and extracellular equations. Based on our earlier findings, hypre is used to solve the linear systems arising from the extracellular equations.

**Table 3:** Execution time (in seconds) for solving the EMI model. We perform 10 time steps of the EMI model with the same setup as in Figure 3 and using hypre to solve the extracellular equations. Measurements are obtained on a 24-core AMD Epyc 7413 CPU, where up to 24 threads are used. The top half of the table shows the initial parallel performance before optimization and parallelization of the right-hand side assembly, whereas the bottom half of the table shows the parallel performance after optimizing and parallelizing the assembly of right-hand sides.

| Threads | Total | Membrane | Intracellular |  | Extracellular |  |
| --- | --- | --- | --- | --- | --- | --- |
|  |  |  | Assembly | Solve | Assembly | Solve |
| 1 | 72.704 | 2.669 | 13.947 | 5.048 | 20.994 | 22.238 |
| 2 | 58.738 | 1.346 | 13.966 | 2.551 | 21.012 | 12.114 |
| 4 | 52.247 | 0.663 | 13.949 | 1.276 | 21.016 | 7.556 |
| 8 | 50.047 | 0.337 | 13.948 | 0.658 | 20.959 | 6.425 |
| 16 | 47.888 | 0.172 | 13.975 | 0.337 | 20.990 | 4.765 |
| 24 | 47.797 | 0.126 | 13.947 | 0.243 | 20.943 | 4.808 |
| 1 | 61.125 | 2.603 | 9.371 | 4.972 | 15.969 | 21.303 |
| 2 | 33.505 | 1.323 | 4.747 | 2.504 | 8.455 | 11.861 |
| 4 | 19.582 | 0.622 | 2.388 | 1.259 | 4.744 | 7.187 |
| 8 | 16.915 | 0.337 | 1.212 | 0.644 | 5.184 | 6.406 |
| 16 | 14.569 | 0.176 | 1.050 | 0.336 | 5.251 | 4.917 |
| 24 | 16.896 | 0.120 | 0.998 | 0.254 | 7.485 | 5.170 |

#### Good speed-up for membrane and intracellular models

Both membrane model and intracellular solve show nearly perfect scalability with speedups of 21.2*×* and 20.8*×*, respectively, when using 24 threads. As we saw earlier, the extracellular solve exhibits a more modest speedup of 4.6*×*. This is expected for iterative solvers of large, sparse linear systems [53], whose performance is limited by memory bandwidth, a resource that is shared by all cores. Solving the intracellular equations does not suffer from this issue due to solving many, small linear systems that are likely to fit within on-chip caches on each CPU core.

#### Inefficient assembly of right-hand sides

In any case, a much bigger concern is posed by the *assembly* of right-hand side vectors for the extracellular and intracellular equations, which takes place during every time step. (Note that left-hand side matrices are assembled only once and their contribution to the overall simulation time is negligible.) In the sequential case, assembly of right-hand sides accounts for half of the overall execution time. Moreover, this part of the computation does not currently benefit from parallel execution, and therefore constitutes a larger share of the overall execution time as more threads are used. Next, we therefore describe code optimizations that aim to parallelize the assembly of right-hand side vectors.

#### Parallel assembly for the intracellular equations

The intracellular linear systems (38)–(40) are all independent, and multiple threads can therefore be used to assemble each of the right-hand sides independently and in parallel. In principle, this is achieved by adding an OpenMP parallel region and worksharing-loop construct (i.e., `#pragma omp parallel for`) to the loop that assembles the intracellular right-hand sides. Unfortunately, this alone is not enough, as we observe severe slowdowns when increasing the number of threads. We traced the cause to the OpenBLAS library, which uses locking to ensure thread-safety in a multithreaded setting. By default, MFEM relies on an external BLAS library to perform factorization and inversion of small, dense matrices during assembly.

To work around the issue, we disable the use of BLAS and fall back to MFEM’s native routines for dense matrix inversion. The result is, first, a performance improvement of 30–50 % from avoiding the overhead of invoking BLAS to invert small matrices. Second, and more importantly, assembly of right-hand sides can now be performed in parallel for the intracellular equations, and it scales almost perfectly up to 8 threads. It is likely that 8 threads are sufficient to saturate the memory bandwidth, and additional threads therefore have little impact on the performance.

#### Parallel assembly for the extracellular equations

Right-hand side assembly for the extracellular equations (41)–(44) is sequential, and limitations in MFEM prevent parallelization for the particular case of using tetrahedral mesh elements. We work around this limitation by adding our own OpenMP-parallel version of the right-hand side assembly, where atomic operations are used to avoid race conditions from multiple threads updating the same element in the right-hand side vector. Ultimately, parallelization speeds up the right-hand side assembly for the extracellular equations by 3.37 times when using 4 threads. Additional threads degrade the performance because of contention and overhead associated with the use of atomic operations.

Due to MFEM being a general finite element framework that supports different kinds of meshes and elements, we observe a non-trivial overhead in its implementation of assembly of right-hand side vectors. In the future, we plan to investigate if a simpler, bespoke code can provide the features that we need while offering further opportunities for optimizing the code.

#### Feynmann optimality

Finally, Table 4 compares the execution time of the parallel EMI solver for tissues of two different sizes. More specifically, we double the size of the tissue to 2.6 mm *×* 1.3 mm and the number of cells to 26*×*65. Roughly speaking, the total execution time also doubles, which indicates that the amount of work performed by the solver is proportional to the tissue area and number of cardiac cells. In [54], Feynman formulated a rule for the complexity of a simulation stating that the computing efforts should only increase linearly with the space-time volume under consideration. For the EMI model, this means that the CPU efforts should only increase linearly with the number of cardiomyocytes and the time interval under consideration. Table 4 indicates that this is the case in these simulations.

**Table 4:** Execution time (in seconds) for solving the EMI model for different meshes. We perform 10 time steps of the EMI model, using hypre to solve the extracellular equations. Measurements are obtained using 24 OpenMP threads on a 24-core AMD Epyc 7413 CPU.

| Cells | Total | Membrane | Intracellular |  | Extracellular |  |
| --- | --- | --- | --- | --- | --- | --- |
|  |  |  | Assembly | Solve | Assembly | Solve |
| 13×65 | 16.90 | 0.12 | 1.00 | 0.25 | 7.49 | 5.17 |
| 26×65 | 34.65 | 0.25 | 2.20 | 0.49 | 13.48 | 12.16 |

## 4 Discussion

Our aim was to provide a comparison of five different models of cardiac electrophysiology in terms of computational efficiency and physiological resolution. We compared the classical bidomain model (BD), the monodomain model (MD), the recently developed Kirchhoff network model (KNM), the associated simplified Kirchhoff network model (SKNM), and the extracellular-membrane-intracellular model (EMI) for a set of classical test cases. The biomarkers used for assessing the properties of the methods are the conduction velocity (CV) and the action potential duration (APD). Both of these biomarkers are well-known to be important characteristics in the analysis of possible arrhythmias, see, e.g., [55, 56, 57, 58, 59] for CVs and [60, 61, 62, 63, 19] for the APDs.

### 4.1 Computational efficiency

The five considered models are different in their basic construction. The classical BD and MD models are developed by averaging over many cardiomyocytes and the models assume that the extracellular space, the intracellular space and the cell membrane all exist everywhere. This represents a colossal simplification of the models in terms of software implementations and computing efforts but clearly limits the models’ validity when the spatial scale is refined towards the size of individual cardiomyocytes. KNM and SKNM are based on representations of individual cells and are as such refining the physiological accuracy compared to BD and MD. Finally, the EMI model is clearly the most detailed one since it represents the cardiomyocytes at the sub-cellular level.

In Figure 2 we compare the spatial and temporal resolutions needed to obtain convergence of the five models. We define convergence to be achieved if the error in the conduction velocity is less than 3%, where the error is computed by comparing the solution obtained by a solution computed with a very fine numerical resolution. The comparison is performed for several values of the cell-to-cell resistance and we note that more resistance requires finer spatial resolution for MD, BD and EMI. However, for KNM and SKNM, the spatial resolution is fixed since it is defined by the cell size; these models do not contain a Δ*x* parameter to be tuned. For EMI the spatial resolution needed is Δ*x* = 5 *µ*m while 10 *µ*m appears to be sufficient for BD and MD. It should be noted that the spatial resolution required for the classical BD and MD models is significantly finer than the meshes frequently used in BD and MD tissue simulations. Standard resolutions appear to be about Δ*x ≈* 0.25 mm, see, e.g., [64, 65, 66, 67, 13]. This means that for 1 cm^3^ cardiac tissue, the number of computational nodes would increase from *∼* 64, 000 in the standard resolution, to one billion in the resolution indicated here. Clearly, this has enormous consequences for the computational demands of the BD and MD models, and we need to emphasize that we do not suggest as a general rule that *∼* 10 *µ*m is necessary for convergence of these models. However, close to infarcted regions, such resolutions appear to be necessary in order to properly compute a converged conduction velocity.

Also, the temporal resolution is estimated using the same criterion. For all models, Δ*t* = 10 *µ*s is sufficient to reach convergence. For the challenging case of simulating conduction in the vicinity of an infarcted region, Figure 3 clearly states that the SKNM method solves the test problem to convergence fastest (3 s) followed by KNM (7 s), MD (4 min), BD (28 min) and EMI (16 h). In Table 3 we showed that the computing efforts of the EMI model could be significantly reduced by parallelizing the code and further improving its performance, but the model is still much more computationally intensive than the other four.

In terms of computational efficiency, SKNM and KNM appear to be very well suited for challenging tissue simulations at small volumes, but the models will be demanding for large volumes since every cardiomyocyte is represented in the model. BD and MD become computationally demanding for challenging tissue simulations but are known to be relevant models, and in fact the only viable option, for large tissues. The EMI model is very accurate but extremely costly in terms of computations and can only be applied when relatively few cells are considered.

### 4.2 Implications for simulating post-infarct arrhythmias

Myocardial infarction induces complex cardiac remodeling such as infiltration of fibrotic tissue [68], up or down regulation of membrane ion channels [69, 70], as well as repositioning of these channels around the cell membrane [71]. While some of these changes can be represented in classical BD/MD formulations through modifications in action potentials and conduction velocities, the finer changes occurring in the cellular scale are lost in these spatially averaged models. This uncertainty has been previously reported in modeling studies using BD/MD formulations which have shown that the inducibility of post infarct reentry is highly sensitive to the conduction velocity parameters chosen to represent the infarct border zone [72, 73]. Figure 4 shows that the faster conduction velocity in the border zone of the MD model results in decreased reentry inducibility as compared to the SKNM model.

Further decreasing the conduction velocity in the border zone resulted in the induction of qualitatively similar reentrant activity in both the MD and SKNM models. Both models resulted in micro-reentrant circuits initiated and maintained within the border zone region. Such complex reentry morphologies have been previously observed in MD simulations that incorporate complex fibrosis within the border zone [74]. While micro-reentrant circuits are difficult to observe experimentally due to limitations in resolution of recording devices, it has been postulated that micro-reentrant circuits could underlie ectopic activity that have been frequently observed in post-infarction hearts [75].

## 5 Conclusion

We have compared five mathematical models of cardiac electrophysiology (BD/MD/KNM/SKNM/EMI) with respect to computational requirements and physiological resolutions. The classical BD and MD models are well suited for simulating large tissues where many cells can be represented by averages. For cell-based accuracy, KNM and SKNM are well suited since every cell is explicitly represented in the simulation. If the impact of varying current density along individual cardiomyocytes needs to be studied, the EMI model is the only viable alternative. On the other hand, the EMI model cannot, at present, be applied to large tissues because of the associated computational load.

## Acknowledgments

This work was partially supported by the European High-Performance Computing Joint Undertaking EuroHPC under grant agreement No. 955495 (MICROCARD) co-funded by the Horizon 2020 programme of the European Union (EU) and the Research Council of Norway.

## Author contributions

K.H.J. and A.T. developed the methodology and designed the experiments. J.D.T. and X.C. enabled and reported the results of the speed up of the EMI model computations, while K.H.J. wrote the remaining simulation code and created the figures. A.T. and H.A. conceived the project. All authors contributed to writing the paper and the final version was approved by all authors.

## Additional information

### Disclosure of writing assistance

The authors used the ChatGPT4 language model to improve the language quality for contributions from non-native English speakers. The authors also rigorously reviewed manuscript to ensure its accuracy and integrity. The authors assume full responsibility for the content of the publication.

## Supplementary Information

### S1 Base model formulation

Here we describe the full base model formulation, taken directly from [38]. In this formulation, the membrane potential (*v*) is given in units of mV, and the Ca^2+^ concentrations are given in units of mM. All currents are expressed in units of A/F, and the Ca^2+^ fluxes are expressed as mmol/ms per total cell volume (i.e., in units of mM/ms). Time is given in ms. The parameters of the model are given in Tables S1–S6.

#### S1.1 The membrane potential

The membrane potential is governed by the equation

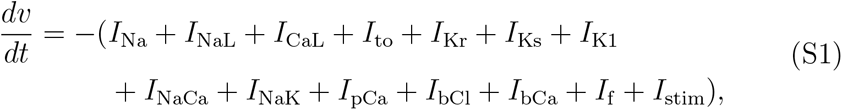

where *I*_stim_ is an applied stimulus current, and *I*_Na_, *I*_NaL_, *I*_CaL_, *I*_to_, *I*_Kr_, *I*_Ks_, *I*_K1_, *I*_NaCa_, *I*_NaK_, *I*_pCa_, *I*_bCl_, *I*_bCa_, and *I*_f_ are membrane currents specified below.

#### S1.2 Stimulation current

In our simulations, *I*_stim_ is given as a constant current of size *−*40 A/F for the 1D simulations and *−*200 A/F for the 2D simulations. The *I*_stim_ current is applied until the membrane potential reaches a value of *−*40 mV. We let the simulation run for 20 ms before applying the stimulation current to allow the model variables to reach a new steady state when parameter changes are applied (i.e., when we include a border zone with an increased extracellular potassium concentration).

#### S1.3 Membrane currents

The currents through the voltage-gated ion channels on the cell membrane are in general given on the form

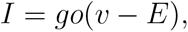

where *g* is the channel conductance, *v* is the membrane potential and *E* is the equilibrium potential of the channel. Furthermore, *o* = Π_*i*_ *z*_*i*_ is the open probability of the channels, where *z*_*i*_ are gating variables, either given as a function of the membrane potential or governed by equations of the form

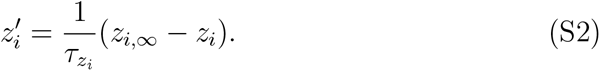

The parameters 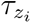 and *z*_*i,∞*_ are specified for each of the gating variables of the model in Table S7.

##### Fast sodium current

The fast sodium current, *I*_Na_, is given by

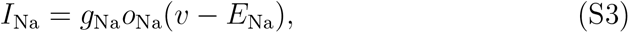

where the open probability is given by

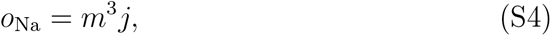

and *m* and *j* are gating variables governed by equations of the form (S2).

##### Late sodium current

The late sodium current, *I*_NaL_, is given by

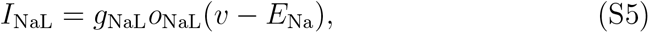

where the open probability is given by

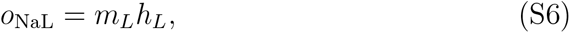

and *m*_*L*_ and *h*_*L*_ are gating variables governed by equations of the form (S2).

##### Transient outward potassium current

The transient outward potassium current, *I*_to_, is given by

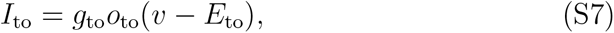

where the open probability is given by

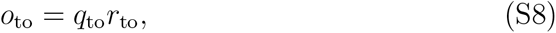

and *q*_to_ and *r*_to_ are gating variables governed by equations of the form (S2).

##### Rapidly activating potassium current

The rapidly activating potassium current, *I*_Kr_, is given by

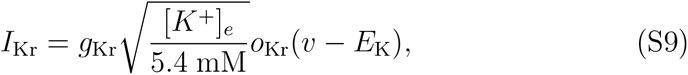

where

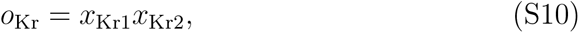

and the dynamics of *x*_Kr1_ and *x*_Kr2_ are governed by equations of the form (S2).

##### Slowly activating potassium current

Theslowly activating potassium current, *I*_Ks_, is given by

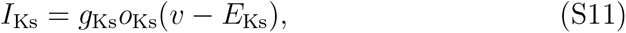

where

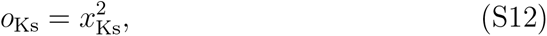

and the dynamics of *x*_Ks_ is governed by an equation of the form (S2).

##### Inward rectifier potassium current

The inward rectifier potassium current, *I*_K1_, is given by

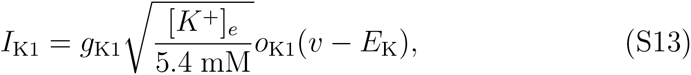

where

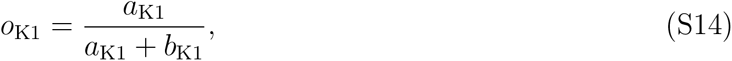

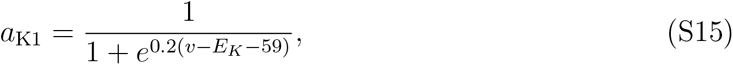

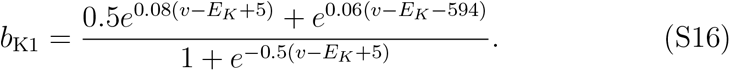

##### Hyperpolarization activated funny current

The hyperpolarization activated funny current, *I*_f_, is given by

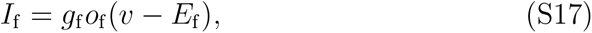

where

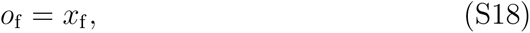

and the dynamics of *x*_f_ is governed by an equation of the form (S2).

##### L-type Ca^2+^ current

The L-type Ca^2+^ current, *I*_CaL_, is given by

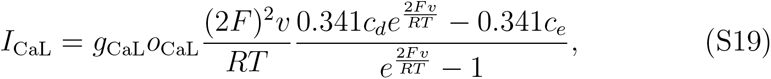

where

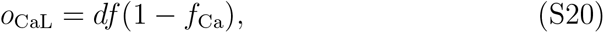

and the dynamics of *d, f* and *f*_Ca_ are governed by equations of the form (S2).

##### Background currents

The background currents, *I*_bCa_ and *I*_bCl_ are given by

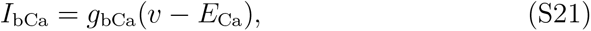

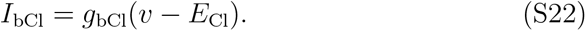

##### Sodium-calcium exchanger

The Na^+^-Ca^2+^ exchanger current, *I*_NaCa_, is given by

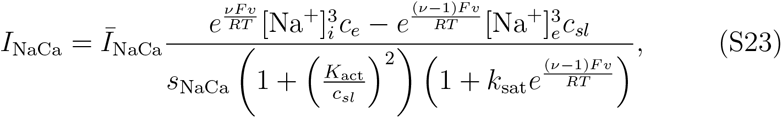

where

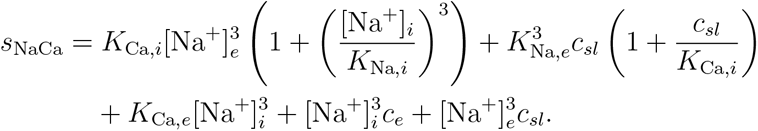

##### Sarcolemmal Ca^2+^ pump

The current through the sarcolemmal Ca^2+^ pump, *I*_pCa_, is given by

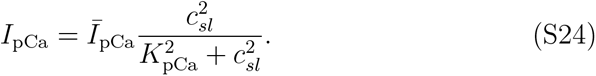

##### Sodium-potassium pump

The current through the Na^+^-K^+^ pump, *I*_NaK_, is given by

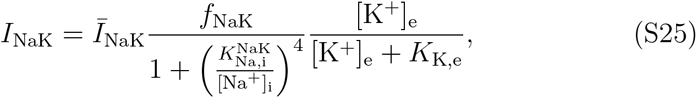

where

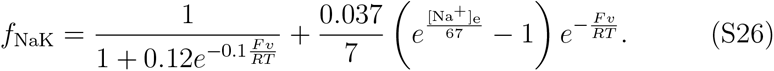

#### S1.4 Ca^2+^ dynamics

The Ca^2+^ dynamics are governed by

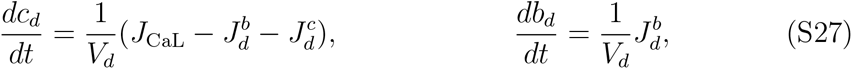

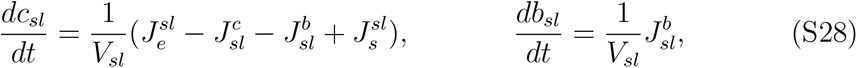

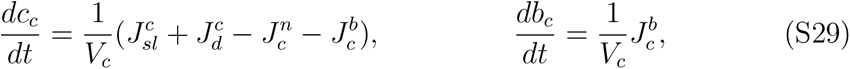

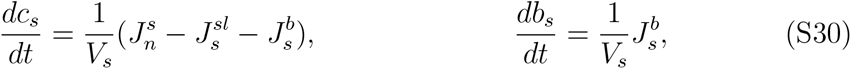

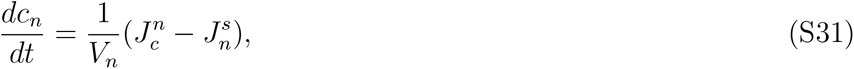

where *c*_*d*_ is the concentration of free Ca^2+^ in the dyad, *b*_*d*_ is the concentration of Ca^2+^ bound to a buffer in the dyad, *c*_*sl*_ is the concentration of free Ca^2+^ in the SL compartment, *b*_*sl*_ is the concentration of Ca^2+^ bound to a buffer in the SL compartment, *c*_*c*_ is the concentration of free Ca^2+^ in the bulk cytosol, *b*_*c*_ is the concentration of Ca^2+^ bound to a buffer in the bulk cytosol, *c*_*s*_ is the concentration of free Ca^2+^ in the jSR, *b*_*s*_ is the concentration of Ca^2+^ bound to a buffer in the jSR, and *c*_*n*_ is the concentration of free Ca^2+^ in the nSR. The expressions for the fluxes are specified below.

#### S1.5 Ca^2+^ fluxes

##### Flux through the SERCA pumps

The flux from the bulk cytosol to the nSR through the SERCA pumps is given by

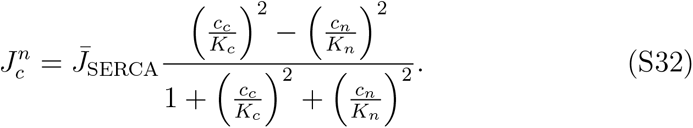

##### Flux through the RyRs

The flux from the jSR to the SL compartment is given by

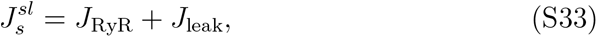

where *J*_RyR_ represents the flux through the active RyR channels and *J*_leak_ represents the flux through the RyR channels that are always open, given by

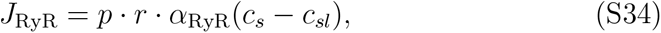

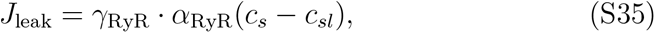

respectively. Here, *p* is the open probability of the active RyR channels given by

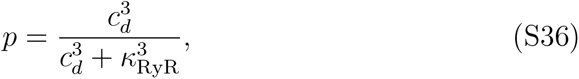

and *r* represents the fraction of RyR channels that are not inactivated and is governed by the equation

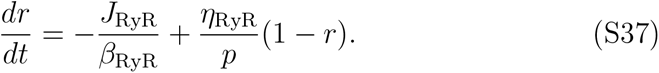

##### Passive diffusion fluxes between compartments

The passive diffusion fluxes between compartments are given by

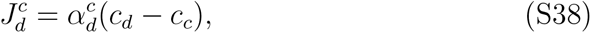

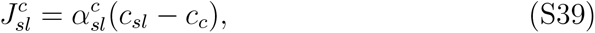

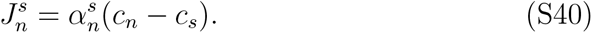

##### Buffer fluxes

The fluxes of free Ca^2+^ binding to a Ca^2+^ buffer are given by

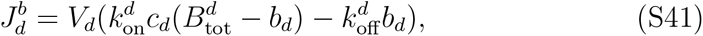

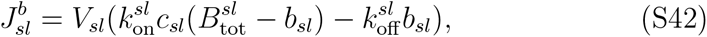

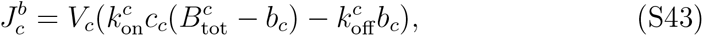

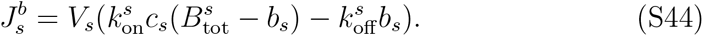

##### Membrane fluxes

The membrane fluxes, *J*_CaL_, *J*_bCa_, *J*_pCa_, and *J*_NaCa_, are given by

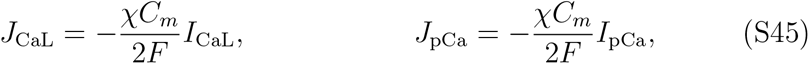

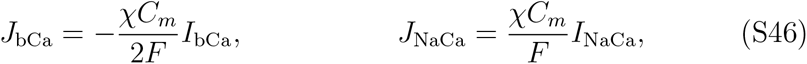

where *I*_CaL_, *I*_bCa_, *I*_pCa_, and *I*_NaCa_ are defined by the expressions given above. Furthermore,

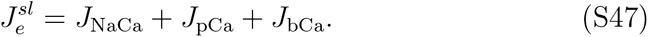

#### S1.6 Nernst equilibrium potentials

The Nernst equilibrium potentials for the ion channels are defined as for the parameter values given in Table S2.

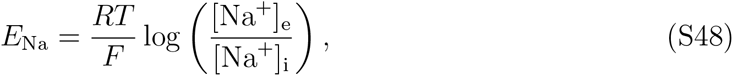

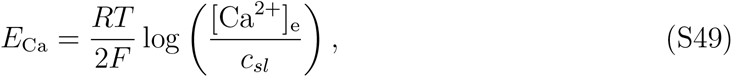

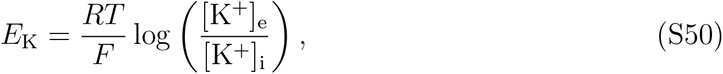

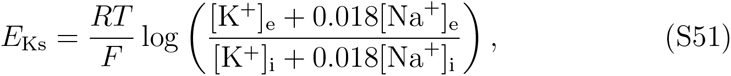

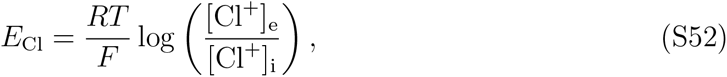

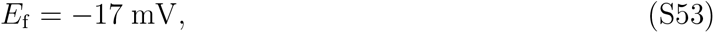

**Table S1:** Default geometry parameters of the base model.

| Parameter | Description | Value |
| --- | --- | --- |
| $V_d$ | Volume fraction of the dyadic subspace | 0.001 |
| $V_{sl}$ | Volume fraction of the SL compartment | 0.028 |
| $V_c$ | Volume fraction of the bulk cytosol | 0.917 |
| $V_s$ | Volume fraction of the jSR | 0.004 |
| $V_n$ | Volume fraction of the nSR | 0.05 |
| $\chi$ | Cell surface to volume ratio | $0.6 \mu\text{m}^{-1}$ |

**Table S2:** Physical constants and ionic concentrations of the base model.

| Parameter | Description | Value |
| --- | --- | --- |
| $C_m$ | Specific membrane capacitance | 0.01 pF/$\mu\text{m}^2$ |
| $F$ | Faraday's constant | 96.485 C/mmol |
| $R$ | Universal gas constant | 8.314 J/(mol·K) |
| $T$ | Temperature | 310 K |
| $[\text{Ca}^{2+}]_e$ | Extracellular $\text{Ca}^{2+}$ concentration | 1.8 mM |
| $[\text{Na}^+]_e$ | Extracellular sodium concentration | 140 mM |
| $[\text{Na}^+]_i$ | Intracellular sodium concentration | 8 mM |
| $[\text{K}^+]_e$ | Extracellular potassium concentration | 5.4 mM |
| $[\text{K}^+]_i$ | Intracellular potassium concentration | 120 mM |
| $[\text{Cl}^-]_e$ | Extracellular chloride concentration | 150 mM |
| $[\text{Cl}^-]_i$ | Intracellular chloride concentration | 15 mM |

**Table S3:**
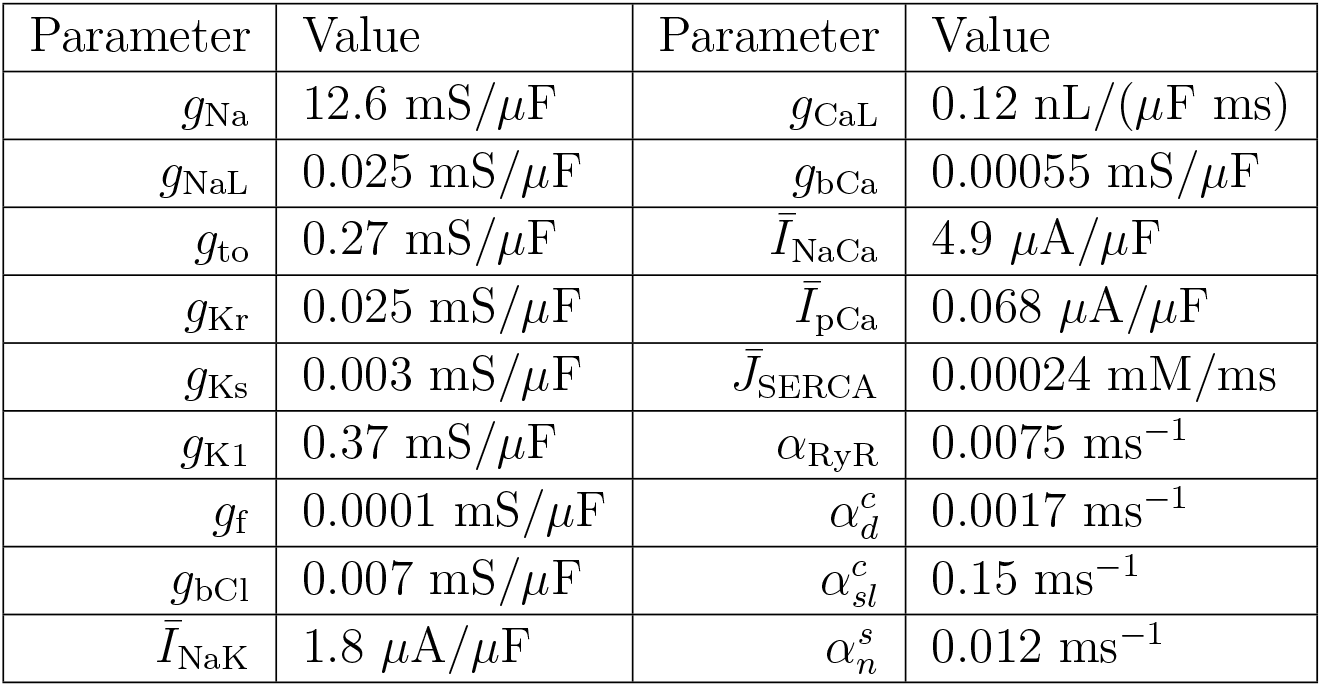
Conductance values and similar parameters for each of the membrane currents and intracellular Ca^2+^ fluxes of the base model.

**Table S4:** Parameters for the intracellular Ca^2+^ fluxes of the base model.

| Parameter | Flux | Value |
| --- | --- | --- |
| $K_c$ | $J_c^n$ | 0.00025 mM |
| $K_n$ | $J_c^n$ | 1.7 mM |
| $\beta_{\text{RyR}}$ | $J_s^{sl}$ | 0.038 mM |
| $\gamma_{\text{RyR}}$ | $J_s^{sl}$ | 0.001 |
| $\kappa_{\text{RyR}}$ | $J_{\text{RyR}}$ | 0.015 mM |
| $\eta_{\text{RyR}}$ | $J_s^{sl}$ | 0.00001 ms<sup>-1</sup> |

**Table S5:** Parameters for the membrane currents of the base model.

| Parameter | Current | Value |
| --- | --- | --- |
| $k_{\text{sat}}$ | $I_{\text{NaCa}}$ | 0.3 |
| $\nu$ | $I_{\text{NaCa}}$ | 0.3 |
| $K_{\text{act}}$ | $I_{\text{NaCa}}$ | 0.00015 mM |
| $K_{\text{Ca},i}$ | $I_{\text{NaCa}}$ | 0.0036 mM |
| $K_{\text{Ca},e}$ | $I_{\text{NaCa}}$ | 1.3 mM |
| $K_{\text{Na},i}$ | $I_{\text{NaCa}}$ | 12.3 mM |
| $K_{\text{Na},e}$ | $I_{\text{NaCa}}$ | 87.5 mM |
| $K_{\text{Na},i}^{\text{NaK}}$ | $I_{\text{NaK}}$ | 11 mM |
| $K_{\text{K},e}$ | $I_{\text{NaK}}$ | 1.5 mM |
| $K_{\text{pCa}}$ | $I_{\text{pCa}}$ | 0.0005 mM |

**Table S6:** Parameters for the Ca^2+^ buffers of the base model.

| Parameter | Compartment | Value |
| --- | --- | --- |
| $B_{\text{tot}}^c$ | Bulk cytosol | 0.07 mM |
| $k_{\text{on}}^c$ | Bulk cytosol | $40 \text{ ms}^{-1}\text{mM}^{-1}$ |
| $k_{\text{off}}^c$ | Bulk cytosol | $0.03 \text{ ms}^{-1}$ |
| $B_{\text{tot}}^d$ | Dyad | 1.2 mM |
| $k_{\text{on}}^d$ | Dyad | $100 \text{ ms}^{-1}\text{mM}^{-1}$ |
| $k_{\text{off}}^d$ | Dyad | $1 \text{ ms}^{-1}$ |
| $B_{\text{tot}}^{sl}$ | Subsarcolemmal space | 0.9 mM |
| $k_{\text{on}}^{sl}$ | Subsarcolemmal space | $100 \text{ ms}^{-1}\text{mM}^{-1}$ |
| $k_{\text{off}}^{sl}$ | Subsarcolemmal space | $0.15 \text{ ms}^{-1}$ |
| $B_{\text{tot}}^s$ | Junctional SR | 27 mM |
| $k_{\text{on}}^s$ | Junctional SR | $100 \text{ ms}^{-1}\text{mM}^{-1}$ |
| $k_{\text{off}}^s$ | Junctional SR | $65 \text{ ms}^{-1}$ |

**Table S7:** Specification of the parameters *z*_*∞*_ and *τ*_*z*_, for *z* = *m, j, m*_*L*_, *h*_*L*_, *d, f, f*_Ca_, *q*_to_, *r*_to_, *x*_Kr1_, *x*_Kr2_, *x*_Ks_ and *x*_f_ in the equations for the gating variables (S2).

| Current | Gate | $z_\infty$ | $\alpha_z$ | $\beta_z$ | $\tau_z$ |
| --- | --- | --- | --- | --- | --- |
| $I_{\text{Na}}$ | $m$ | $\frac{1}{(1 + e^{(-57-v)/9})^2}$ | $0.13e^{-((v+46)/16)^2}$ | $0.06e^{-((v-5)/51)^2}$ | $\alpha_m + \beta_m$ |
| | $j$ | $\frac{1}{(1 + e^{(v+72)/7})^2}$ | $\begin{cases} 0, & \text{if } v \geq -40 \\ \frac{\begin{pmatrix} -2.5 \cdot 10^4 e^{0.2v} \\ -7 \cdot 10^{-6} e^{-0.04v} \end{pmatrix} (v+38)}{1 + e^{0.3(v+79)}}, & \text{otherwise} \end{cases}$ | $\begin{cases} \frac{0.6e^{0.06v}}{1 + e^{-0.1(v+32)}}, & \text{if } v \geq -40 \\ \frac{0.02e^{-0.01v}}{1 + e^{-0.14(v+40)}}, & \text{otherwise} \end{cases}$ | $\frac{1}{\alpha_j + \beta_j}$ |
| $I_{\text{NaL}}$ | $m_L$ | $\frac{1}{1 + e^{(-43-v)/5}}$ | $\frac{1}{6.8e^{(v+12)/35}}$ | $8.6e^{-(v+77)/6}$ | $\alpha_m + \beta_m$ |
| | $h_L$ | $\frac{1}{1 + e^{(v+88)/7.5}}$ | | | 200 ms |
| $I_{\text{CaL}}$ | $d$ | $\frac{1}{1 + e^{-(v+5)/6}}$ | $\frac{1 - e^{-\frac{v+5}{6}}}{0.035(v+5)}$ | | $\alpha_d d_\infty$ |
| | $f$ | $\frac{1}{1 + e^{(v+35)/9}} + \frac{0.6}{1 + e^{(50-v)/20}}$ | $\frac{1}{0.02e^{-(0.034(v+14.5)^2)} + 0.02}$ | | $\alpha_f$ |
| | $f_{\text{Ca}}$ | $\frac{1.7c_d}{1.7c_d + 0.012}$ | $\frac{1}{1.7c_d + 0.012}$ | | $\alpha_{\text{Ca}}$ |
| $I_{\text{to}}$ | $q_{\text{to}}$ | $\frac{1}{1 + e^{(v+53)/13}}$ | $\frac{39}{0.57e^{-0.08(v+44)} + 0.065e^{0.1(v+46)}}$ | 6 | $\alpha_{q_{\text{to}}} + \beta_{q_{\text{to}}}$ |
| | $r_{\text{to}}$ | $\frac{1}{1 + e^{-(v-22.3)/18.75}}$ | $\frac{14.4}{e^{0.09(v+30.61)} + 0.37e^{-0.12(v+24)}}$ | 2.75 | $\alpha_{r_{\text{to}}} + \beta_{r_{\text{to}}}$ |
| $I_{\text{Kr}}$ | $x_{\text{Kr1}}$ | $\frac{1}{1 + e^{-(v+20.7)/4.9}}$ | $\frac{450}{1 + e^{-(v+45)/10}}$ | $\frac{6}{1 + e^{(v+30)/11.5}}$ | $\alpha_{x_{\text{Kr1}}} \cdot \beta_{x_{\text{Kr1}}}$ |
| | $x_{\text{Kr2}}$ | $\frac{1}{1 + e^{(v+88)/50}}$ | $\frac{3}{1 + e^{-(v+60)/20}}$ | $\frac{1.12}{1 + e^{(v-60)/20}}$ | $\alpha_{x_{\text{Kr2}}} \cdot \beta_{x_{\text{Kr2}}}$ |
| $I_{\text{Ks}}$ | $x_{\text{Ks}}$ | $\frac{1}{1 + e^{-(v+3.8)/14}}$ | $\frac{990}{1 + e^{-(v+2.4)/14}}$ | | $\alpha_{x_{\text{Ks}}}$ |
| $I_{\text{f}}$ | $x_{\text{f}}$ | $\frac{1}{1 + e^{(v+78)/5}}$ | $\frac{1900}{1 + e^{(v+15)/10}}$ | | $\alpha_{x_{\text{Ks}}}$ |

#### S2 Definition of conduction velocity

In the 1D strand simulations for computing the conduction velocity, the first cell of the cell strand (or a corresponding length) was stimulated. The conduction velocity was computed as the distance between the centers of cells 3 and 13 divided by the duration of time between when the membrane potential in these two points reached a value above *−*20 mV.

#### S3 MFEM configuration

The numerical experiments presented in Section 3.6 use MFEM 4.5.2, which is configured with the following options:

- `CMAKE BUILD TYPE=Release`
- `MFEM USE MPI=YES`
- `MFEM USE OPENMP=YES`
- MFEM THREAD SAFE=YES
- `MFEM USE LAPACK=NO`
- `MFEM USE METIS=YES` and `MFEM USE METIS 5=YES`
- `MFEM USE HYPRE=YES`
- `MFEM USE SUITESPARSE=YES`
- `MFEM USE SUPERLU=YES`

We note especially that the option MFEM THREAD SAFE=YES is required to ensure thread-safe versions of certain MFEM functions are used. This is necessary due to assembling and solving multiple linear systems in parallel. In addition, the use of BLAS to invert small, dense matrices during assembly of right-hand side vectors is disabled with the option `MFEM USE LAPACK=NO`. Besides MFEM, several additional software packages are needed, including METIS 5.1.0, hypre 2.26.0, SuiteSparse 5.12.0, SuperLU dist 8.1.0 and OpenMPI 4.1.4. We further note that OpenMP support is enabled in hypre (i.e., `--with-openmp`) and SuperLU dist (i.e., `enable openmp=ON`). The code is compiled using GCC 11.4.0.

Finally, the listing below shows the parallel code that was used for assembling right-hand side vector of the extracellular equations. The code is based on the function `LinearForm::Assemble()` from MFEM. It is some-what simplified because only boundary integrals are required for our particular case. A parallel region and worksharing-loop construct were added, as well as atomic operations to avoid race conditions when updating the entries of the right-hand side vector. Furthermore, we note that calls to the function `Mesh::GetBdrElementTransformation()` were substituted with a thread-safe version to avoid race conditions and thus ensure correct results.

~~~
void linearform_assemble_omp ( LinearForm & b)
{
    b = 0.0;
    */* Only boundary integrals are needed in our case. */*
    Array < LinearForm Integrator * > * boundary_integs = b. GetBLFI ();
    if ( boundary_integs - > Size ())
    {
        Finite ElementSpace * fes = b. FESpace (); Mesh * mesh = fes - > GetMesh ();
        # pragma omp parallel for
        for ( int i = 0; i < fes -> GetNBE (); i++)
        {
            Array <int > vdofs;
            IsoparametricTransformation eltrans;
            DofTransformation * doftrans;
            Vector elemvect;
            mesh -> GetBdrElementTransformation (i, & eltrans );
            for ( int k =0; k < boundary_integs - > Size (); k ++)
            {
                (* boundary_integs )[ k]- > Assemble RHSElementVect (
                       * fes - > GetBE ( i), eltrans , elemvect );
                double * bx = b. HostRead Write ();
                double * elemvectx = elemvect. HostRead Write ();
                for ( int l = 0; l < vdofs. Size (); l++) {
                      int j = vdofs[ l];
                      if ( j >= 0) {
                           # pragma omp atomic
                           bx[ j] += elemvectx [ l];
                      } else {
                           # pragma omp atomic
                           bx[-1 - j] -= elemvectx [ l];
                      }
                }
            }
        }
    }
}
~~~

## References

[1] Yoram Rudy and Jonathan R Silva. Computational biology in the study of cardiac ion channels and cell electrophysiology. Quarterly Reviews of Biophysics, 39(1):57–116, 2006.

[2] Bogdan Amuzescu, Razvan Airini, Florin Bogdan Epureanu, Stefan A Mann, Thomas Knott, and Beatrice Mihaela Radu. Evolution of mathematical models of cardiomyocyte electrophysiology. Mathematical Biosciences, 334:108567, 2021.

[3] Piero C Franzone, Luca F Pavarino, and Simone Scacchi. Mathematical Cardiac Electrophysiology, volume 13. Springer, 2014.

[4] Karoline Horgmo Jæger and Aslak Tveito. Differential equations for studies in computational electrophysiology. Simula SpringerBriefs on Computing, 2023.

[5] JC Neu and W Krassowska. Homogenization of syncytial tissues. Critical Reviews in Biomedical Engineering, 21(2):137–199, 1993.

[6] Craig S Henriquez and Wenjun Ying. The bidomain model of cardiac tissue: from microscale to macroscale. Cardiac Bioelectric Therapy: Mechanisms and Practical Implications, pages 211–223, 2021.

[7] Karoline H Jæger and Aslak Tveito. Deriving the bidomain model of cardiac electrophysiology from a cell-based model; properties and comparisons. Frontiers in Physiology, page 2439, 2022.

[8] Microcard. https://www.microcard.eu.

[9] Piero Colli Franzone and Giuseppe Savaré. Degenerate evolution systems modeling the cardiac electric field at micro-and macroscopic level. In Evolution Equations, Semigroups and Functional Analysis: in memory of Brunello Terreni, pages 49–78. Springer, 2002.

[10] Aslak Tveito, Karoline H Jæger, Miroslav Kuchta, Kent-Andre Mardal, and Marie E Rognes. A cell-based framework for numerical modeling of electrical conduction in cardiac tissue. Frontiers in Physics, 5:48, 2017.

[11] Aslak Tveito, Karoline H Jæger, Glenn T Lines,Lukasz Paszkowski, Joakim Sundnes, Andrew G Edwards, Tuomo Māki-Marttunen, Geir Halnes, and Gaute T Einevoll. An evaluation of the accuracy of classical models for computing the membrane potential and extracellular potential for neurons. Frontiers in Computational Neuroscience, 11:27, 2017.

[12] Karoline H Jæger and Aslak Tveito. Derivation of a cell-based mathematical model of excitable cells. In Modeling Excitable Tissue, pages 1–13. Springer, Cham, 2020.

[13] Karoline H Jæger, Andrew G Edwards, Wayne R Giles, and Aslak Tveito. From millimeters to micrometers; re-introducing myocytes in models of cardiac electrophysiology. Frontiers in Physiology, 12:763584, 2021.

[14] Ada J Ellingsrud, Andreas Solbrå, Gaute T Einevoll, Geir Halnes, and Marie E Rognes. Finite element simulation of ionic electrodiffusion in cellular geometries. Frontiers in Neuroinformatics, 14:11, 2020.

[15] Ada J Ellingsrud, Cécile Daversin-Catty, and Marie E Rognes. A cell-based model for ionic electrodiffusion in excitable tissue. In Modeling Excitable Tissue, pages 14–27. Springer, Cham, 2021.

[16] Åshild Telle, James D Trotter, Xing Cai, Henrik Finsberg, Miroslav Kuchta, Joakim Sundnes, and Samuel T Wall. A cell-based framework for modeling cardiac mechanics. Biomechanics and Modeling in Mechanobiology, pages 1–25, 2023.

[17] Giacomo Rosilho de Souza, Simone Pezzuto, and Rolf Krause. Effect of gap junction distribution, size, and shape on the conduction velocity in a cell-by-cell model for electrophysiology. In Olivier Bernard, Patrick Clarysse, Nicolas Duchateau, Jacques Ohayon, and Magalie Viallon, editors, Functional Imaging and Modeling of the Heart, volume 13958 of Lecture Notes in Computer Science, pages 117–126, Cham, June 2023. Springer.

[18] Karoline H Jæger, Andrew G Edwards, Andrew McCulloch, and Aslak Tveito. Properties of cardiac conduction in a cell-based computational model. PLoS Computational Biology, 15(5):e1007042, 2019.

[19] Karoline H Jæger, Andrew G Edwards, Wayne R Giles, and Aslak Tveito. Arrhythmogenic influence of mutations in a myocyte-based computational model of the pulmonary vein sleeve. Scientific Reports, 12(1):1–18, 2022.

[20] Alessio Paolo Buccino, Miroslav Kuchta, Karoline Horgmo Jæger, Torbjørn Vefferstad Ness, Pierre Berthet, Kent-Andre Mardal, Gert Cauwenberghs, and Aslak Tveito. How does the presence of neural probes affect extracellular potentials? Journal of Neural Engineering, 16(2):026030, 2019.

[21] Kristian G Hustad, Ena Ivanovic, Adrian L Recha, and Abinaya Abbi Sakthivel. Conduction velocity in cardiac tissue as function of ion channel conductance and distribution. In Computational Physiology: Simula Summer School 2021-Student Reports, pages 41–50. Springer International Publishing Cham, 2022.

[22] Joshua Steyer, Fatemeh Chegini, Mark Potse, Axel Loewe, and Martin Weiser. Continuity of microscopic cardiac conduction in a computational cell-by-cell model. In Computing in Cardiology, volume 50, Atlanta, Georgia, USA, October 2023.

[23] Karoline H Jæger, Kristian Gregorius Hustad, Xing Cai, and Aslak Tveito. Efficient numerical solution of the EMI model representing the extracellular space (E), cell membrane (M) and intracellular space (I) of a collection of cardiac cells. Frontiers in Physics, 8:539, 2021.

[24] Giacomo Rosilho de Souza, Rolf Krause, and Simone Pezzuto. Boundary integral formulation of the cell-by-cell model of cardiac electrophysiology. Engineering Analysis with Boundary Elements, 158:239–251, 2024.

[25] Pietro Benedusi, Paola Ferrari, Marie Rognes, and Stefano Serra-Capizzano. Modeling excitable cells with the EMI equations: spectral analysis and iterative solution strategy. Journal of Scientific Computing, 2024.

[26] Nanna Berre, Marie E Rognes, and Andre Massing. Cut finite element discretizations of cell-by-cell EMI electrophysiology models. arXiv preprint arXiv:2306.03001, 2023.

[27] Fernando Henríquez, Carlos Jerez-Hanckes, and Fernando Altermatt. Boundary integral formulation and semi-implicit scheme coupling for modeling cells under electrical stimulation. Numerische Mathematik, 136(1):101–145, 2017.

[28] Ngoc Mai Monica Huynh, Fatemeh Chegini, Luca Franco Pavarino, Martin Weiser, and Simone Scacchi. Convergence analysis of BDDC preconditioners for hybrid DG discretizations of the cardiac cell-by-cell model. SIAM Journal on Scientific Computing, 45(6):A2836–A2857, 2023.

[29] Fakhrielddine Bader, Mostafa Bendahmane, Mazen Saad, and Raafat Talhouk. Microscopic tridomain model of electrical activity in the heart with dynamical gap junctions. part 2 – derivation of the macroscopic tridomain model by unfolding homogenization method. Asymptotic Analysis, 132(3-4):575–606, apr 2023.

[30] Fakhrielddine Bader, Mostafa Bendahmane, Mazen Saad, and Raafat Talhouk. Microscopic tridomain model of electrical activity in the heart with dynamical gap junctions. Part 1–modeling and well-posedness. Acta Applicandae Mathematicae, 179(1):11, 2022.

[31] Joyce Reimer, Sebastián A Domínguez-Rivera, Joakim Sundnes, and Raymond J Spiteri. Physiological accuracy in simulating refractory cardiac tissue: the volume-averaged bidomain model vs. the cell-based EMI model. bioRxiv, pages 2023–04, 2023.

[32] Fatemeh Chegini, Algiane Froehly, Ngoc Mai Monica Huynh, Luca F Pavarino, Mark Potse, Simone Scacchi, and Martin Weiser. Efficient numerical methods for simulating cardiac electrophysiology with cellular resolution. In 10th International Conference on Computational Methods for Coupled Problems in Science and Engineering, 2023.

[33] Mark Potse, Luca Cirrottola, and Algiane Froehly. A practical algorithm to build geometric models of cardiac muscle structure. In 8th European Congress on Computational Methods in Applied Sciences and Engineering (ECCOMAS), Oslo, Norway, June 2022.

[34] Karoline Horgmo Jæger and Aslak Tveito. Efficient, cell-based simulations of cardiac electrophysiology; the Kirchhoff network model (KNM). NPJ Systems Biology and Applications, 9(1):25, 2023.

[35] Karoline Horgmo Jæger and Aslak Tveito. The simplified Kirchhoff network model (SKNM): a cell-based reaction–diffusion model of excitable tissue. Scientific Reports, 13(1):16434, 2023.

[36] NHL Kuijpers, RH Keldermann, T Arts, and PAJ Hilbers. Computer simulations of successful defibrillation in decoupled and non-uniform cardiac tissue. EP Europace, 7(2):S166–S177, 2005.

[37] Jan P Kucera, Stephan Rohr, and Yoram Rudy. Localization of sodium channels in intercalated disks modulates cardiac conduction. Circulation Research, 91(12):1176–1182, 2002.

[38] Karoline H Jæger, Verena Charwat, Bérénice Charrez, Henrik Finsberg, Mary M Maleckar, Sam Wall, Kevin Healy, and Aslak Tveito. Improved computational identification of drug response using optical measurements of human stem cell derived cardiomyocytes in microphysiological systems. Frontiers in Pharmacology, 10:1648, 2020.

[39] Joakim Sundnes, Glenn Terje Lines, and Aslak Tveito. An operator splitting method for solving the bidomain equations coupled to a volume conductor model for the torso. Mathematical Biosciences, 194(2):233–248, 2005.

[40] Stanley Rush and Hugh Larsen. A practical algorithm for solving dynamic membrane equations. IEEE Transactions on Biomedical Engineering, 4:389–392, 1978.

[41] Johan Hake, Henrik Finsberg, Kristian Gregorius Hustad, and George Bahij. Gotran – General ODE TRANslator, 2020. https://github.com/ComputationalPhysiology/gotran.

[42] Leonardo Dagum and Ramesh Menon. OpenMP: An industry-standard API for shared-memory programming. IEEE Computational Science and Engineering, 5(1):46–55, 1998.

[43] Robert Anderson, Julian Andrej, Andrew Barker, Jamie Bramwell, Jean-Sylvain Camier, Jakub Cerveny, Veselin Dobrev, Yohann Dudouit, Aaron Fisher, Tzanio Kolev, Will Pazner, Mark Stowell, Vladimir Tomov, Ido Akkerman, Johann Dahm, David Medina, and Stefano Zampini. MFEM: A modular finite element methods library. Computers & Mathematics with Applications, 81:42–74, 2021.

[44] Robert Anderson, Julian Andrej, Andrew Barker, Jamie Bramwell, Jean-Sylvain Camier, Jakub Cerveny, Veselin Dobrev, Yohann Dudouit, Aaron Fisher, Tzanio Kolev, et al. MFEM: A modular finite element methods library. Computers & Mathematics with Applications, 81:42–74, 2021.

[45] Joachim Berdal Haga, Harald Osnes, and Hans Petter Langtangen. Efficient block preconditioners for the coupled equations of pressure and deformation in highly discontinuous media. International Journal for Numerical and Analytical Methods in Geomechanics, 35(13):1466–1482, 2011.

[46] Christoph Geuzaine and Jean-François Remacle. Gmsh: a threedimensional finite element mesh generator with built-in pre- and post-processing facilities. International Journal for Numerical Methods in Engineering, 79:1309–1331, 2009.

[47] Karoline H Jæger, Kristian Gregorius Hustad, Xing Cai, and Aslak Tveito. Operator splitting and finite difference schemes for solving the EMI model. In Modeling Excitable Tissue, pages 44–55. Springer, Cham, 2020.

[48] hypre: High performance preconditioners. https://llnl.gov/casc/hypre, https://github.com/hypre-space/hypre.

[49] Timothy A. Davis. Algorithm 832: UMFPACK v4.3—an unsymmetric-pattern multifrontal method. ACM Trans. Math. Softw., 30(2):196–199, 2004.

[50] Xiaoye S. Li and James W. Demmel. SuperLU DIST: A scalable distributed-memory sparse direct solver for unsymmetric linear systems. ACM Trans. Math. Softw., 29(2):110–140, 2003.

[51] Madison S Spach, J Francis Heidlage, Paul C Dolber, and Roger C Barr. Mechanism of origin of conduction disturbances in aging human atrial bundles: experimental and model study. Heart Rhythm, 4(2):175–185, 2007.

[52] Nele Vandersickel, Ivan V Kazbanov, Anita Nuitermans, Louis D Weise, Rahul Pandit, and Alexander V Panfilov. A study of early afterdepolarizations in a model for human ventricular tissue. PloS one, 9(1):e84595, 2014.

[53] Hartwig Anzt, Erik Boman, Rob Falgout, Pieter Ghysels, Michael Heroux, Xiaoye Li, Lois Curfman McInnes, Richard Tran Mills, Sivasankaran Rajamanickam, Karl Rupp, Barry Smith, Ichitaro Yamazaki, and Ulrike Meier Yang. Preparing sparse solvers for exascale computing. Philosophical Transactions of the Royal Society A: Mathematical, Physical and Engineering Sciences, 378(2166):20190053, 2020.

[54] Richard P Feynman. Simulating physics with computers. International Journal of Theoretical Physics, 21(6):467–488, 1982.

[55] Joel Kupersmith, Ehud Krongrad, and Albert L Waldo. Conduction intervals and conduction velocity in the human cardiac conduction system: studies during open-heart surgery. Circulation, 47(4):776–785, 1973.

[56] Bo Han, Mark L Trew, and Callum M Zgierski-Johnston. Cardiac conduction velocity, remodeling and arrhythmogenesis. Cells, 10(11):2923, 2021.

[57] James H King, Christopher L-H Huang, and James A Fraser. Determinants of myocardial conduction velocity: implications for arrhythmogenesis. Frontiers in Physiology, 4:154, 2013.

[58] Edward Vigmond, Caroline Roney, Jason D Bayer, and Kumaraswamy Nanthakumar. The accuracy of cardiac surface conduction velocity measurements. medRxiv, pages 2024–01, 2024.

[59] Patrick M Boyle, William H Franceschi, Marion Constantin, Claudia Hawks, Thomas Desplantez, Natalia A Trayanova, and Edward J Vigmond. New insights on the cardiac safety factor: Unraveling the relationship between conduction velocity and robustness of propagation. Journal of Molecular and Cellular Cardiology, 128:117–128, 2019.

[60] Paul F Cranefield. Action potentials, afterpotentials, and arrhythmias. Circulation Research, 41(4):415–423, 1977.

[61] Isabelle Banville and Richard A Gray. Effect of action potential duration and conduction velocity restitution and their spatial dispersion on alternans and the stability of arrhythmias. Journal of Cardiovascular Electrophysiology, 13(11):1141–1149, 2002.

[62] Chien-Suu Kuo, K Munakata, C Pratap Reddy, and B Surawicz. Characteristics and possible mechanism of ventricular arrhythmia dependent on the dispersion of action potential durations. Circulation, 67(6):1356–1367, 1983.

[63] Karoline Horgmo Jæger, Andrew G Edwards, Wayne R Giles, and Aslak Tveito. A computational method for identifying an optimal combination of existing drugs to repair the action potentials of SQT1 ventricular myocytes. PLoS Computational Biology, 17(8):e1009233, 2021.

[64] Steven Niederer, Lawrence Mitchell, Nicolas Smith, and Gernot Plank. Simulating human cardiac electrophysiology on clinical time-scales. Frontiers in Physiology, 2:14, 2011.

[65] Steven A Niederer, Eric Kerfoot, Alan P Benson, Miguel O Bernabeu, Olivier Bernus, Chris Bradley, Elizabeth M Cherry, Richard Clayton, Flavio H Fenton, Alan Garny, et al. Verification of cardiac tissue electrophysiology simulators using an n-version benchmark. Philosophical Transactions of the Royal Society A: Mathematical, Physical and Engineering Sciences, 369(1954):4331–4351, 2011.

[66] Richard H Clayton and Alexander V Panfilov. A guide to modelling cardiac electrical activity in anatomically detailed ventricles. Progress in Biophysics and Molecular Biology, 96(1-3):19–43, 2008.

[67] Fagen Xie, Zhilin Qu, Junzhong Yang, Ali Baher, James N Weiss, Alan Garfinkel, et al. A simulation study of the effects of cardiac anatomy in ventricular fibrillation. The Journal of Clinical Investigation, 113(5):686–693, 2004.

[68] Virpi Talman and Heikki Ruskoaho. Cardiac fibrosis in myocardial infarction – from repair and remodeling to regeneration. Cell and Tissue Research, 365(3):563–581, 2016.

[69] Pradeep S. Rajendran, Keijiro Nakamura, Olujimi A. Ajijola, Marmar Vaseghi, J. Andrew Armour, Jeffrey L. Ardell, and Kalyanam Shivkumar. Myocardial infarction induces structural and functional remodelling of the intrinsic cardiac nervous system. The Journal of Physiology, 594(2):321–341, 2016.

[70] M. Amoni, E. Dries, S. Ingelaere, D. Vermoortele, H.L. Roderick, P. Claus, R. Willems, and K.R. Sipido. Ventricular arrhythmias in ischemic cardiomyopathy—new avenues for mechanism-guided treatment. Cells, 10(10):2629, 2021.

[71] W.E. Louch, H.K. Mørk, J. Sexton, T.A. Strømme, P. Laake, I. Sjaastad, and O.M. Sejersted. T-tubule disorganization and reduced synchrony of Ca2+ release in murine cardiomyocytes following myocardial infarction. Journal of Physiology, 574:519–533, 2006.

[72] C. Mendonca Costa, G. Plank, C.A. Rinaldi, S.A. Niederer, and M.J. Bishop. Modeling the electrophysiological properties of the infarct border zone. Frontiers in Physiology, 9:356, 2018.

[73] P. Colli-Franzone, V. Gionti, L.F. Pavarino, S. Scacchi, and C. Storti. Role of infarct scar dimensions, border zone repolarization properties and anisotropy in the origin and maintenance of cardiac reentry. Mathematical Biosciences, 315:108228, 2019.

[74] Rafael Sachetto Oliveira, Sergio Alonso, Fernando Otaviano Campos, Bernardo Martins Rocha, João Filipe Fernandes, Titus Kuehne, and Rodrigo Weber Dos Santos. Ectopic beats arise from micro-reentries near infarct regions in simulations of a patient-specific heart model. Scientific Reports, 8(1):16392, 2018.

[75] Hector Martinez-Navarro, Ana Mincholé, Alfonso Bueno-Orovio, and Blanca Rodriguez. High arrhythmic risk in antero-septal acute myocardial ischemia is explained by increased transmural reentry occurrence. Scientific Reports, 9(1):16803, 2019.

